# A novel virus-induced cyclic dinucleotide, 2′3′-c-di-GMP, mediates STING-dependent antiviral immunity in *Drosophila*

**DOI:** 10.1101/2023.05.08.539652

**Authors:** Hua Cai, Lihua Li, Kailey Slavik, Jingxian Huang, Ting Yin, Léna Hédelin, Zhangmin Xiang, Yunyun Yang, Xiaoyan Li, Yuqiang Chen, Ziming Wei, Huimin Deng, Di Chen, Renjie Jiao, Nelson Martins, Carine Meignin, Philip Kranzusch, Jean-Luc Imler

## Abstract

In mammals, the enzyme cGAS senses the presence of cytosolic DNA and synthesizes the cyclic dinucleotide (CDN) 2′3′-cGAMP. This CDN binds to and activates the protein STING to trigger immunity. We recently discovered in the model organism *Drosophila melanogaster* two cGAS-like receptors (cGLRs) that activate STING-dependent antiviral immunity and can produce 3′2′-cGAMP, in addition to 2′3′-cGAMP. Here we explore CDN-mediated immunity in 14 different *Drosophila* species covering 50 million years of evolution and report that 2′3′-cGAMP and 3′2′-cGAMP fail to control infection by *Drosophila* C virus in *D. serrata, D. sechellia* and *D. mojavensis*. Using an accurate and sensitive mass spectrometry method, we discover an unexpected diversity of CDNs produced in a cGLR-dependent manner in response to viral infection in *D. melanogaster*, including a novel CDN, 2′3′-c-di-GMP. We show that 2′3′-c-di-GMP is the most potent STING agonist identified so far in *D. melanogaster* and that this molecule also activates a strong antiviral transcriptional response in *D. serrata*. Our results shed light on the evolution of cGLRs in flies and provide a basis for the understanding of the function and regulation of this emerging family of PRRs in animal innate immunity.

## Introduction

Innate immunity is the first line of host-defense against infections. It involves a set of pattern recognition receptors (PRRs), which sense conserved microbial products and activate signaling to mount an efficient antimicrobial response. The model organism of the fruit fly *Drosophila melanogaster* has played an important role in the identification of the first characterized family of PRRs, the Toll-like receptors^1–3^. In *Drosophila*, however, Toll functions as a receptor for the cytokine Spaetzle, rather than as a PRR^4^. Sensing of bacterial and fungal infection occurs upstream of Toll, and is mediated by members of two families of PRRs, peptidoglycan recognition proteins (PGRPs) and Gram-negative binding proteins (GNBPs), which detect lysine-type peptidoglycan or β-glucans^5–7^. Other members of the PGRP family sense DAP-type peptidoglycan to activate the IMD pathway^8–11^.

Viral infections have long been thought to be controlled solely by RNA interference in insects, even though induction of gene expression in virus-infected flies has been noted (reviewed in^12, 13^). A transcriptional response to viral infection involving the evolutionarily conserved protein STING has indeed been recently characterized in flies and silkworms^14–16^. In mammals, STING is activated upon binding a second messenger, the cyclic dinucleotide (CDN) 2′3′-cGAMP, and triggers the transcription factors IRF3 and NF-*K*B to induce expression of genes encoding interferons and other antiviral molecules (reviewed in^17^). 2′3′-cGAMP is produced by the enzyme cyclic GMP-AMP synthase (cGAS), which acts as a PRR sensing the presence of double stranded (ds)DNA in the cytosol^18^. In flies and silkworms, STING regulates Relish, a member of the NF-*K*B family, to induce expression of genes that function in concert to control of viral infections^14, 15, 19^. The pathway is activated by two cGAS-like receptors (cGLRs), which, together with cGAS, define a new family of PRRs in *Drosophila*^20, 21^.

Interestingly, cGAS-STING signaling originated in bacteria, where it plays a critical role in the control of phage infections (reviewed in^22^). A large collection of cGAS/DncV-like nucleotidyltransferases (CD-NTases) are activated upon phage infection and catalyze the production of cyclic di-or trinucleotides as part of a cyclic oligonucleotide-based antiphage signaling system (CBASS)^23, 24^. These phage-induced cyclic oligonucleotides bind to a variety of effector proteins, including homologs of STING, activating them to oppose phage replication, either by triggering cell death or by degrading phage nucleic acids^25, 26^. Thus, it appears that animals acquired the components of the cGAS-STING pathway from prokaryotes, a hypothesis supported by the recent demonstration that STING signaling is activated and participates in the response to infection in invertebrates beyond insects, e.g., the sea anemone *Nematostella vectensis*^27^, and even in the choanoflagellate *Monosiga brevicollis*, a unicellular close relative of metazoans^28^.

In *D. melanogaster*, two cGLRs have been shown to produce CDNs that activate STING signaling and their initial characterization revealed interesting differences compared to mammalian cGAS^20, 21^. First, the activity of cGLR1 can be triggered *in vitro* or in transfected cells by dsRNA rather than dsDNA. The ligand activating cGLR2 is still unknown, although mutation of conserved residues critical for DNA or RNA binding in cGAS and oligoadenylate synthetase decrease the activity of cGLR2, suggesting allosteric activation by a nucleic acid. Second, although cGLR2 also produces 2′3′-cGAMP, both cGLRs were found to produce a novel potent STING agonist, the CDN 3′2′-cGAMP. Third, *Drosophila* genomes encode a remarkable number of putative cGLRs, with between 2 and 7 cGAS homologues per species. Of note, cGLR1 and cGLR2 have so far mostly been molecularly characterized *in vitro* with recombinant proteins or using transfected cells. Here, we exploit the biodiversity of the *Drosophila* genus to gain insight on the contribution of 2′3′-cGAMP and 3′2′-cGAMP in the control of viral infection and on the function of cGLRs *in vivo*. Our results reveal a rapidly evolving family of PRRs which participate in host-defense through the production of diverse nucleotide signals.

## Results

### 3′2′-cGAMP induces antiviral immunity in most, but not all, Drosophila species

To investigate the role of 3′2′-cGAMP signaling in antiviral immunity in the *Drosophila* genus, we selected 14 species covering 50 million years of evolution (Fig. 1A). We injected them with 3′2′-cGAMP and 2′3′-cGAMP, the ancestral signaling molecule in metazoans, as well as 2′3′-c-di-AMP, identified as a minor product of cGLR1^20, 21^. The flies were subsequently challenged with *Drosophila* C virus (DCV, *Dicistroviridae* family), a natural *Drosophila* pathogen that is sensitive to CDN signaling in *D. melanogaster*, and viral RNA load was monitored two-and three-days post-infection (Fig. 1B). As with 3′2′-cGAMP and 2′3′-cGAMP, injection of 2′3′-c-di-AMP resulted in significant reduction of DCV RNA in *D. melanogaster*, revealing that the production by this CDN *in vitro* by recombinant cGLR1 might be physiologically relevant (Fig. 1C)^20, 21^. Injection of the three CDNs also resulted in decreased viral RNA loads in 8 other fly species (Fig. 1B,D; Suppl. Fig. 1). However, neither 2′3′-cGAMP nor 2′3′-c-di-AMP reduced DCV load in *D. sechellia*, *D. serrata*, *D. kikkawai*, *D. pseudoobscura* and *D. mojavensis* (Fig. 1B,E,F; Suppl. Fig. 1). The injection of 3′2′-cGAMP led to reduced DCV replication in 11 of the 14 species, including *D. melanogaster*, and in most cases the decrease in viral RNAs was stronger than in flies injected with 2′3′-c-di-AMP or 2′3′-cGAMP. Yet, intriguingly, neither 3′2′-cGAMP nor the other tested CDNs affected DCV RNA levels in *D. sechellia*, *D. serrata* or *D. mojavensis* (Fig. 1B,F; Suppl. Fig. 1). Overall, these results confirm the importance of 3′2′-cGAMP in antiviral immunity in *Drosophila* and raise the possibility that other types of CDNs mediate antiviral protection in some *Drosophila* species that respond poorly to 3′2′-cGAMP, 2′3′-cGAMP and 2′3′-c-di-AMP.

**Figure 1:**
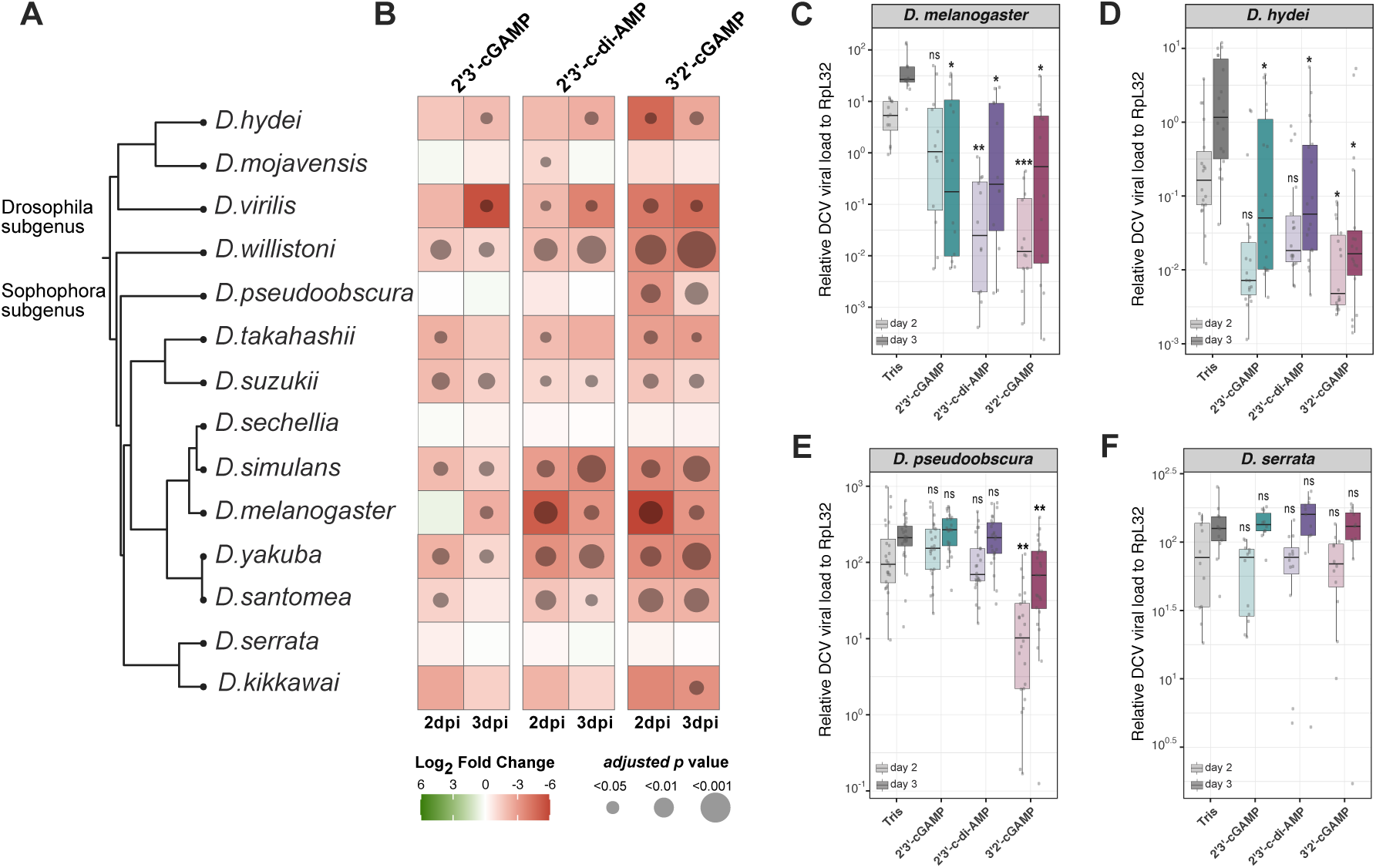
3′2′-cGAMP induces antiviral immunity in most, but not all, *Drosophila* species. **A.** Phylogeny of the 14 *Drosophila* species used. Drosophila and Sophophora subgenus are indicated. **B.** Summary of the antiviral effect of the indicated CDNs in the *Drosophila* species. Heatmap showing the log2 fold change of DCV RNA load at two-and three-days post-infection of flies that had been pre-injected with the indicated CDNs or Tris. Significant changes (adjusted *p* value ≤0.05; pairwise permutation test with FDR method) are highlighted with dots. **C-F.** Relative DCV RNA loads at two or three days post DCV infection in flies from the indicated species. Data are from at least three independent experiments, each performed in biological triplicates and shown with boxplot with scatter plot. Analysis using pairwise permutation test with FDR method upon CDN-compared to Tris-injection is shown (ns= non significant, *<0.05, **<0.01 and ***<0.001).

### cGLR1 and cGLR2 produce several CDNs in vivo

To detect CDNs produced by *D. melanogaster* cGLR1 and cGLR2 *in vivo*, we took advantage of transgenic fly lines. We previously reported that the STING pathway is activated in the absence of viral infection in flies expressing wild-type cGLR1 or 2, but not catalytically inactive mutant versions, suggesting that CDNs are produced^20^. To monitor the presence of CDNs, we designed a bioassay based on the injection of hemolymph from cGLR-overexpressing flies into naïve flies, followed by analysis of expression of *STING-regulated genes* (*srg*)^19^ (Suppl. Fig. 2A). Injection of hemolymph extracted from flies ectopically expressing cGLR1, but not the inactive cGLR1^AFA^ mutant, into wild-type flies resulted in significant upregulation of *srg2* and *srg3* expression (Suppl. Fig. 2D,E). Similar trends for increased expression were observed for the genes *Drosophila STING* (*dSTING*) itself and *srg1*. Comparable results were observed for the transfer of hemolymph from flies ectopically expressing cGLR2. No such induction was observed when the hemolymph was transferred into *dSTING* mutant flies (Suppl. Fig. 2B–E). Overall, these results indicate that a STING agonist is produced and circulating in the cGLR-overexpressing flies.

We next developed a liquid chromatography tandem mass spectrometry (LC-MS/MS) method for detection and quantification of CDNs in the hemolymph. We also optimized the extraction procedure, the purification and the solid phase separation to detect CDNs in fly lysates (Suppl. Fig. 3, 4, 5)^20, 29^. Both 2′3′-cGAMP and 3′2′-cGAMP were detected in the hemolymph of flies ectopically expressing cGLR1, but not the catalytically inactive version (Fig. 2A; Suppl. Fig. 4A,B,D,E). These CDNs were also present in the hemolymph of the flies ectopically expressing cGLR2. Interestingly, two other CDNs were detected in the hemolymph of cGLR2 transgenic flies, 2′3′-c-di-AMP and a previously undescribed CDN, 2′3′-c-di-GMP (Fig. 2A; Suppl. Fig. 4C,F). None of these CDNs were present in the hemolymph of flies expressing the catalytically inactive version of cGLR2. 2′3′-c-di-GMP, in addition to 2′3′-cGAMP, 3′2′-cGAMP, and 2′3′-c-di-AMP, was also present in extracts prepared from lysates of flies ectopically expressing catalytically active cGLR2 (Fig. 2B). Note that 2′3′-c-di-AMP, together with some 3′3′-c-di-AMP, was also detected in control flies expressing the inactive enzyme or GFP. To rule out a contribution of the microbiota, we analyzed extracts from human HEK293T cells transfected with a cGLR2-expressing vector and observed the production of 3′2′-cGAMP, 2′3′-cGAMP, 2′3′-c-di-GMP and, albeit in lower quantities, 2′3′-c-di-AMP (Fig. 2C; Suppl. Fig. 6). In summary, we have established a method to detect CDNs *in vivo* in *Drosophila*. This revealed the production of 2′3′-c-di-GMP, which has until now only been characterized as a synthetic analog of 3′3′-c-di-GMP^30^.

**Figure 2:**
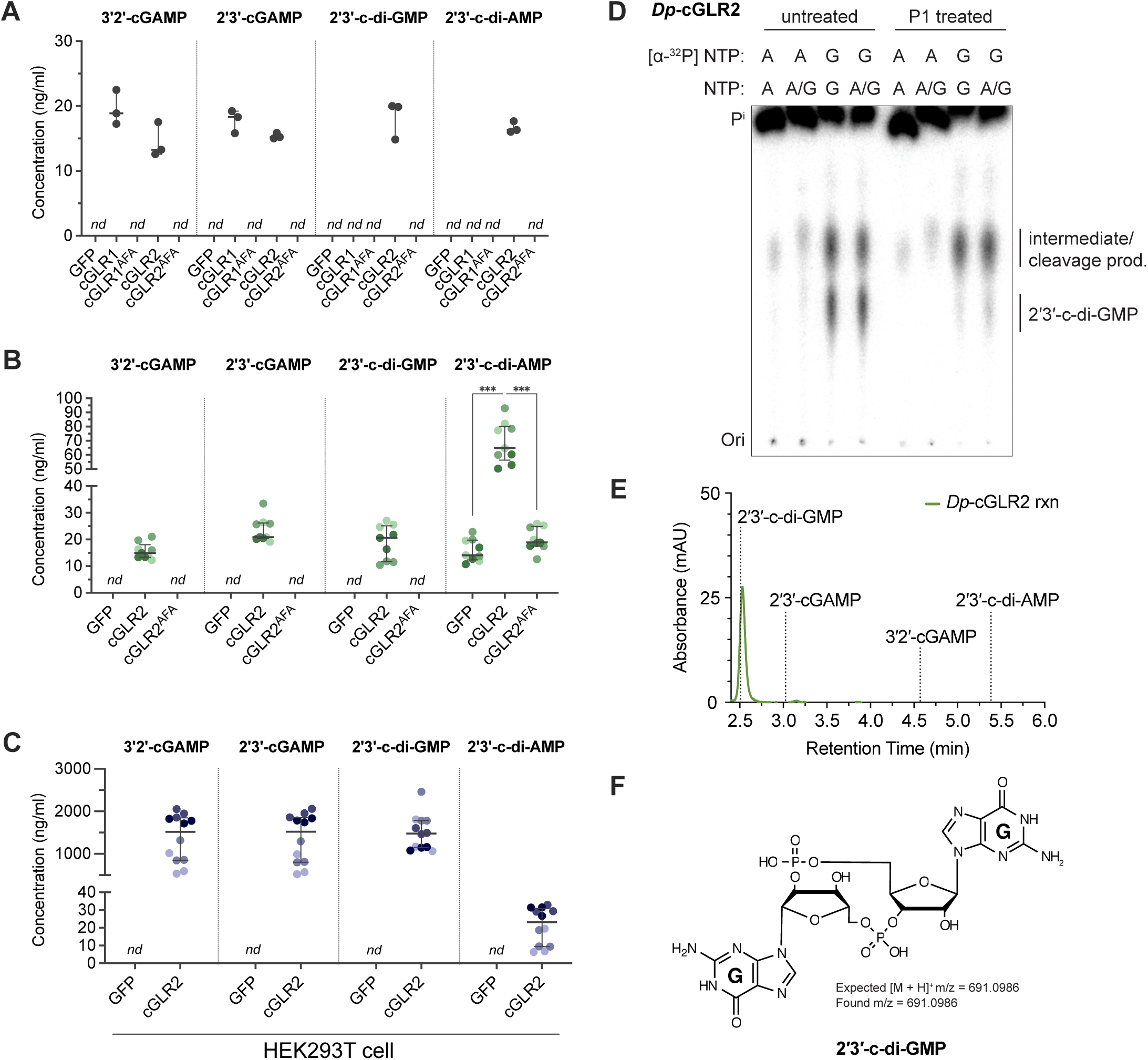
cGLR1 and cGLR2 produce several CDNs in *D. melanogaster* flies. **A-B.** Concentration of CDNs measured by LC-MS in the hemolymph (**A**) or in whole fly lysates (**B**) collected from transgenic flies ectopically expressing cGLR1, cGLR2, their inactive cGLR^AFA^ versions or a control transgene (GFP). **C.** Concentration of CDNs measured by LC-MS in the lysate of HEK293T cells overexpressing cGLR2 or GFP. Data are representative of one (A), three (B) or four (C) independent experiments and shown with dot plots with median and quartile. *nd*, non-detected. Concentration of 2’3’-c-di-AMP in whole flies was analysed by pairwise permutation test with FDR method (***<0.001). **D.** Thin-layer chromatography (TLC) analysis of α-^32^P labeled *Dp*-cGLR2 reaction products and treatment with P1 nuclease, which cleaves 3′–5′ phosphodiester linkages. Specific labeling reveals that only G nucleobases are incorporated in the cGLR2 product and P1 treatment supports that the cGLR2 product contains one 3′–5′ bond and one P1-insensitive bond. Data are representative of n= 3 independent experiments. **E.** HPLC chromatogram showing C18 elution profile of *Dp*-cGLR2 reaction; dashed lined indicates retention times of synthetic nucleotide standards. Data are representative of n= 3 independent experiments. **F.** High-resolution mass spectrometry confirms the major *Dp*-cGLR2 product as 2′3′-c-di-GMP. See Fig. S4, S5, S6, S7G,H.

**Figure 3:**
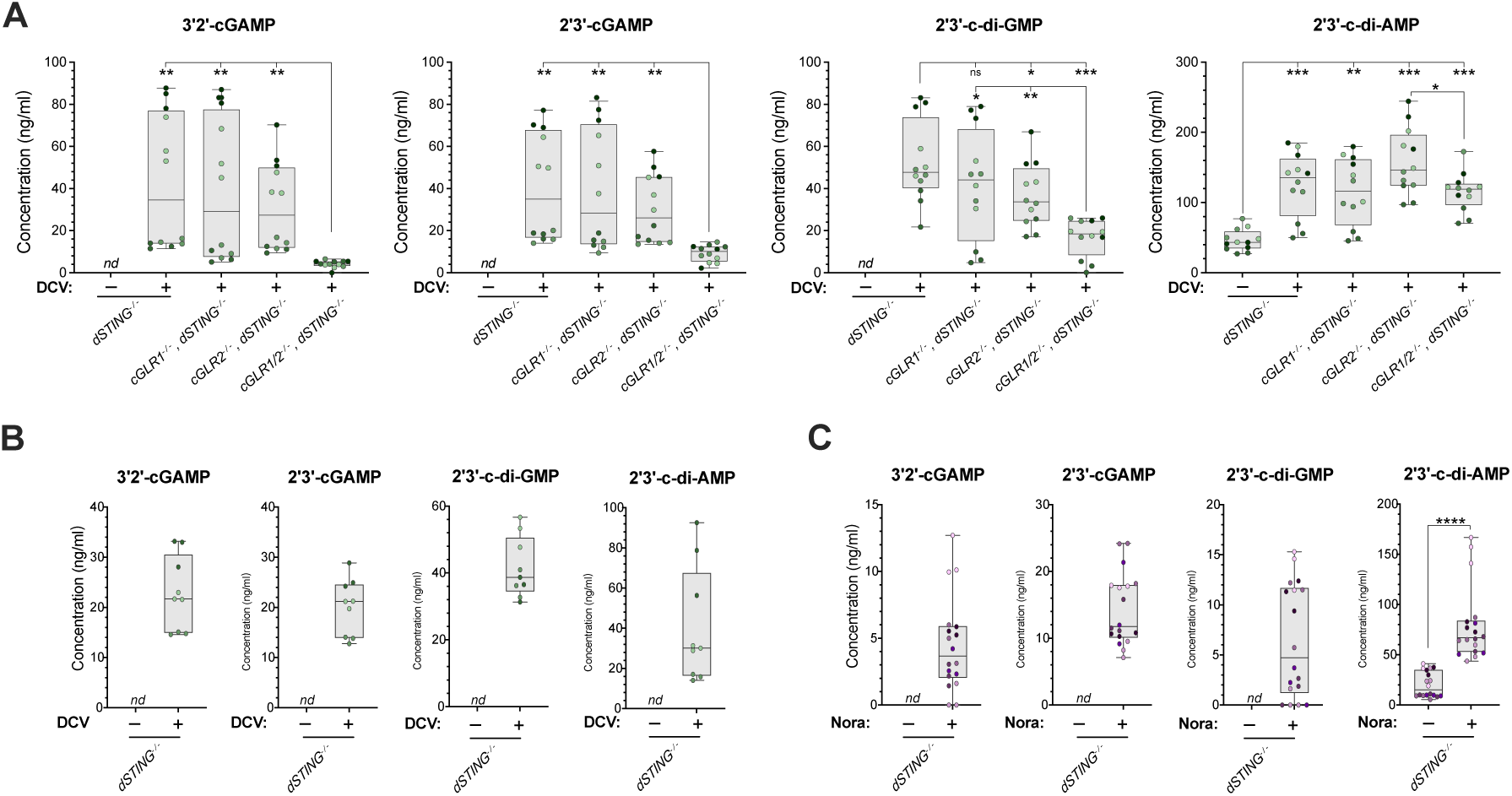
DCV infection triggers cGLR-dependent production of CDNs in *D. melanogaster*. **A.** Concentration of CDNs measured by LC-MS in whole fly lysates from Tris-or DCV-injected flies. Data are from at least four independent experiments. **B.** Concentration of CDNs in the hemolymph of Tris-or DCV-injected *dSTING* knockout flies. Data are from three independent experiments. Each experiment involves 1000 male flies. **C.** Concentration of CDNs in whole fly lysates from Nora-infected *dSTING* knockout flies. Data are from four independent experiments, each involving 500 male flies. All data are shown with boxplot with scatter plot and were analysed using permutation test with FDR method (ns= non significant, *<0.05, **<0.01, ***<0.001 and ****<0.0001), non-detected (*nd*).

**Figure 4:**
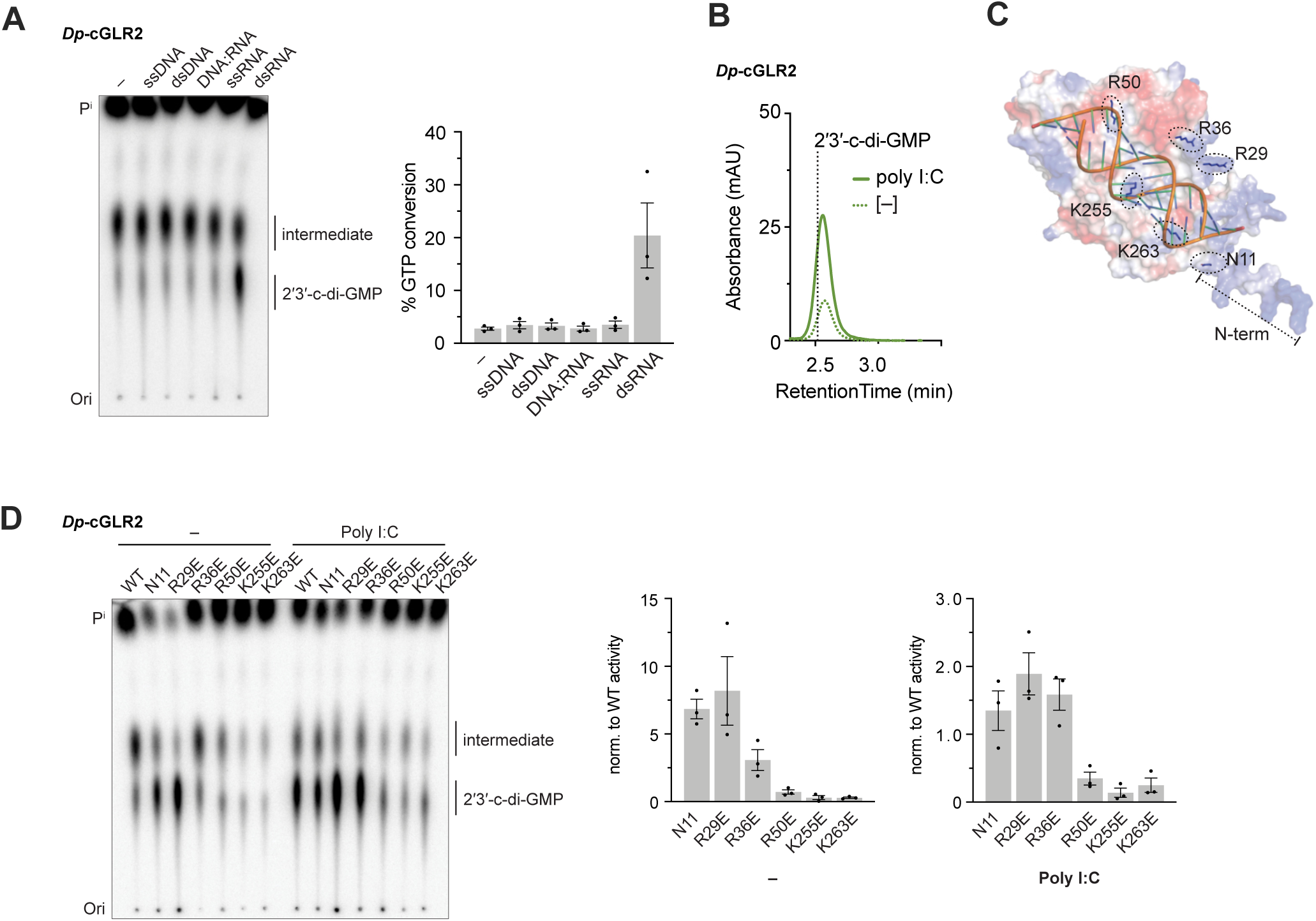
dsRNA binding of Drosophila cGLR2 facilitates its production of 2’3’-c-di-GMP. **A.** TLC analysis and quantification of GTP conversion to 2′3′-c-di-GMP by *Dp*-cGLR2 in the presence of different nucleic acid ligands. Data are mean +/-s.e.m. of n= 3 individual experiments. **B.** HPLC chromatogram showing 2′3′-c-di-GMP synthesis by *Dp*-cGLR2 in the presence of dsRNA mimic Poly I:C or a buffer control. Data are representative of n= 3 independent experiments. **C.** Surface electrostatic view of predicted *Dp*-cGLR2 structure modeled with a 19 bp dsRNA ligand, highlighting predicted interacting residues selected for mutagenesis. **D.** TLC analysis and quantification of wildtype (WT) and mutant *Dp*-cGLR2 activity in the absence and presence of dsRNA mimic Poly I:C. For quantification, % GTP conversion to 2′3′-c-di-GMP for each reaction was normalized to the wildtype control mean value for either buffer or Poly I:C reactions. Data are mean +/-s.e.m. of n= 3 individual experiments.

**Figure 5:**
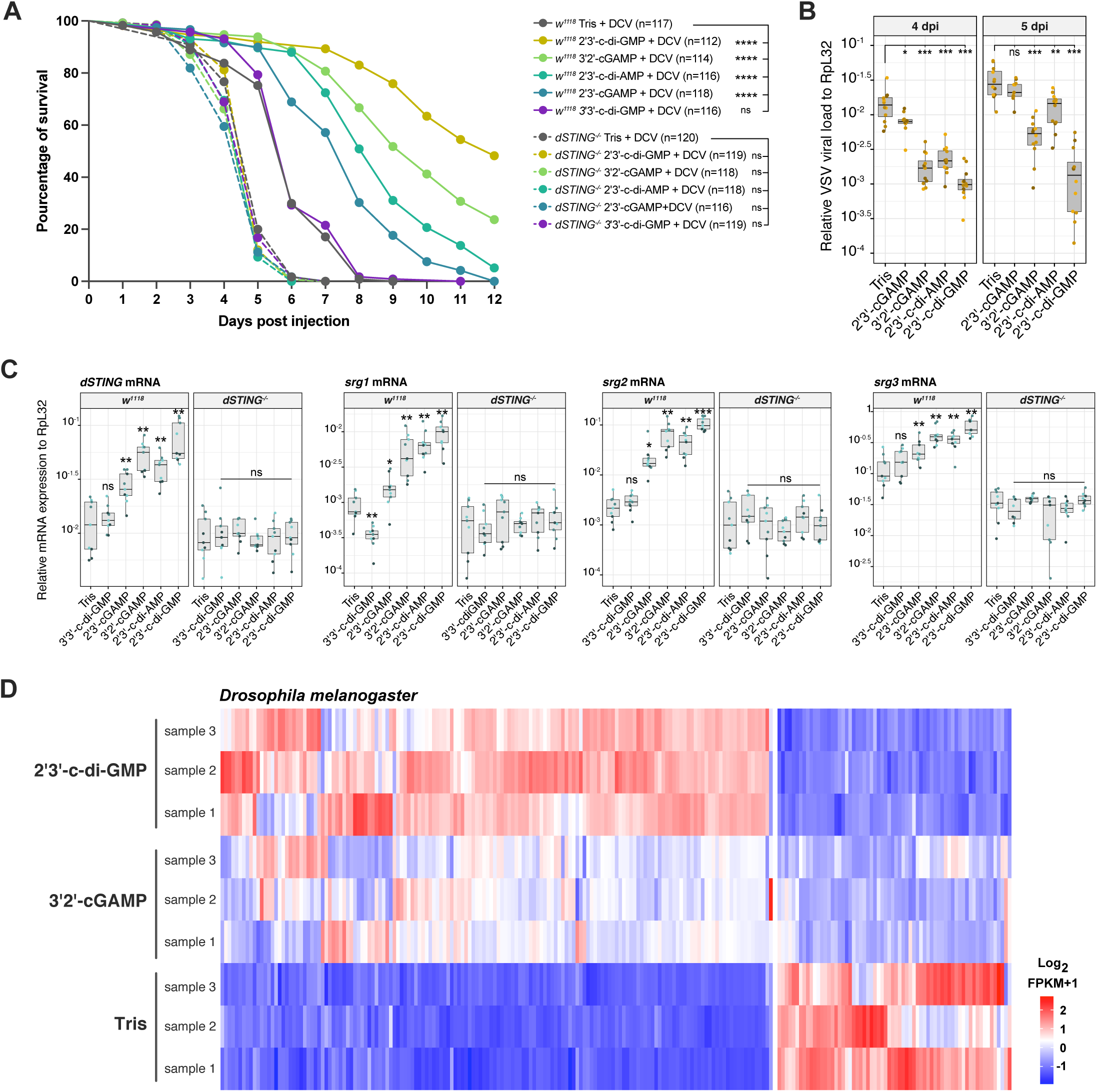
2’3’-c-di-GMP triggers a potent antiviral response in *D. melanogaster*. **A.** Survival of control *(isogenic w*^1118^) and *dSTING* knockout flies pretreated with the indicated CDNs before systemic DCV infection. Data are from 4 independent experiments, each with 3 independent groups of around 10 flies and analysed by Log-rank test (ns= non significant and ****<0.0001). **B.** Control flies pre-injected with the indicated CDNs or Tris after 7 days were challenged with VSV and viral RNA load was monitored after 4-or 5-days post injection. Data are from four independent experiments, each performed in biological triplicates (n = 12). Data are shown with boxplot with scatter plot and were analysed using permutation test with FDR method (ns= non significant, *<0.05, **<0.01, ***<0.001 and ****<0.0001). **C.** Relative gene expression of the *STING-regulated genes dSTING*, *srg1*, *srg2* and *srg3* 7 days after injection of the indicated CDNs in control and *dSTING* knockout flies. Samples were collected from three independent experiments, each involving three groups of 6 flies including 3 males and 3 females. Data are shown with boxplot with scatter plot and were analysed using permutation test with FDR method (ns= non significant, *<0.05, **<0.01 and ***<0.001). **D.** Expression profiles of control *D. melanogaster (isogenic w*^1118^) 7 days after injection with Tris, 3′2′-cGAMP or 2′3′-c-di-GMP. All differentially expressed genes (DEGs) including upregulated and downregulated genes are shown in log2(FPKM+1). Data are from three independent experiments.

In our experiments the production of 2′3′-c-di-GMP in flies correlated exclusively with the expression of catalytically active cGLR2 (Fig. 2A,B). To determine if cGLR2 produces 2′3′-c-di-GMP as a nucleotide second messenger, we investigated the *in vitro* activity of this enzyme. We first performed a systematic screen of cGLR2 homologs encoded in 13 *Drosophila* species (Suppl. Fig. S7A,E) and discovered that *D. pseudoananassae* (*Dp*) cGLR2 and *D. bipectinata* (*Db*) cGLR2 synthesize CDN products *in vitro* (Fig. 2D; S7E). Targeted mutation of the *Dp*-cGLR2 metal-coordinating residue D82 confirmed that CDN synthesis was dependent on the conserved cGLR catalytic site (Fig. S8B)^31^. Using nucleobase specific labeling and nuclease digestion to analyze the cGLR2 product, we determined that this CDN is produced exclusively from GTP substrates and contains a P1-insensitive phosphodiester linkage (Fig. 2D; Fig. S7E) indicating the presence of a 2′–5′ bond. We next analyzed cGLR2 reactions using high-performance liquid chromatography (HPLC) and observed that the major cGLR2 product exhibits a C18 chromatography profile identical to a 2′3′-c-di-GMP synthetic standard (Fig. 2E; Fig. S7D,F). Using LC-MS/MS analysis in comparison to the synthetic standard, we confirmed that 2′3′-c-di-GMP is major product of *Drosophila* cGLR2 (Fig. S7G,H). Interestingly, we observed that cGLR2 exclusively synthesizes 2′3′-c-di-GMP, in contrast to the production of both 3′2′-cGAMP and 2′3′-c-di-AMP by cGLR1^21^. We note that both cGLR1 and cGLR2 exhibit more specific CDN synthesis activity *in vitro* than upon overexpression in flies.

### DCV infection triggers cGLR-dependent production of CDNs in Drosophila

We next monitored production of CDNs in virus-infected flies. We used *dSTING* mutant flies in these experiments to maximize the ability to detect free CDNs^32^. Only 2′3′-c-di-AMP could be detected in the extracts from non-infected flies (note that 3′3′-c-di-AMP was also present in the extracts). However, systemic DCV infection resulted in the induction of 2′3′-cGAMP, 3′2′-cGAMP and 2′3′-c-di-GMP. The quantity of 2′3′-c-di-AMP also significantly increased in DCV-infected flies (Fig. 3A). In addition, 2′3′-cGAMP, 3′2′-cGAMP, 2′3′-c-di-GMP and 2′3′-c-di-AMP were detected in the hemolymph of DCV-infected *dSTING* mutant flies (Fig. 3B). Expression of the four CDNs was also upregulated following enteric infection with another natural *Drosophila* pathogen, Nora virus^33^ (Fig. 3C). To assess the contribution of cGLRs to this response, we repeated the experiment using *cGLR1/2* mutant flies recombined in *dSTING* null mutant background. Production of 2′3′-cGAMP and 3′2′-cGAMP in response to DCV infection decreased significantly when both cGLR1 and cGLR2 were absent but was not affected in single mutant flies (Fig. 3A). A small but significant decrease of 2′3′-c-di-GMP was observed in *cGLR2* mutant flies. The concentration of 2′3′-c-di-GMP further dropped when both *cGLR1* and *cGLR2* were mutated. Intriguingly, we note that production of these three CDNs could still be detected in the *cGLR1/2* double mutant flies. By contrast with 2′3′-cGAMP, 3′2′-cGAMP and 2′3′-c-di-GMP, the upregulation of 2′3′-c-di-AMP by DCV infection was still observed in the absence of cGLR1 and cGLR2 (Fig. 3A).

### dsRNA binding of Drosophila cGLR2 facilitates its production of 2′3′-c-di-GMP

We previously determined that *Drosophila* cGLR1 functions as a dsRNA sensor, consistent with its role in CDN production during RNA virus infection^20, 21^. To determine how cGLR2 is activated to produce CDNs, we monitored *in vitro* CDN synthesis in the presence of different nucleic acid ligands (Fig. 4A,B). We found that *Dp*-and *Db*-cGLR2 synthesize a low amount of 2′3′-c-di-GMP in the absence of exogenous nucleic acid ligands and that activation is enhanced most strongly in the presence of dsRNA (Fig. 4A, Fig. S8A). We additionally used HPLC to monitor *Dp*-cGLR2 reactions and verified that dsRNA facilitates 2′3′-c-di-GMP synthesis (Fig. 4B).

To understand how cGLR2 recognizes nucleic acid ligands we modeled *Dp*-cGLR2 with a short dsRNA ligand and designed charge-swap mutations to conserved basic residues in the predicted interaction surface (Fig. 4C). Many of these residues are conserved in cGAS and *Drosophila* cGLR1, providing a structural explanation for the role of cGLR2 as a nucleic acid sensor^21, 34–36^ (Fig. S7B). In order to understand the role of the highly basic, disordered N-terminus of *Dp-*cGLR2, we additionally generated a truncation mutant starting at residue N11. *Dp*-cGLR2 truncation and charge-swap mutants variably disrupted CDN synthesis by cGLR2 (Fig. 4D). Surprisingly, truncation of the *Dp*-cGLR2 N-terminus resulted in ligand-independent nucleotide product synthesis, suggesting that the disordered N-terminus negatively regulates enzymatic activity in the absence of dsRNA. A charge swap mutation to R29 similarly resulted in robust, ligand-independent product synthesis by cGLR2. Similar to cGAS and *Drosophila* cGLR1, basic residues R50, K255, and K263 are critical for cGLR2 activity, supporting a common role for these positions in nucleic acid sensing by cGLR enzymes^21, 34–36^. Together our mutational analysis of *Dp*-cGLR2 supports that the shared cGLR ligand binding surface is critical for enzymatic activity of cGLR2 and that conserved basic residues control dsRNA-induced CDN synthesis.

### 2′3′-c-di-GMP triggers a potent antiviral response in flies

We previously reported that injection of 2′3′-cGAMP and 3′2′-cGAMP, but not 3′3′-c-di-GMP, primes antiviral immunity, such that injected flies display significantly increased resistance to DCV^19, 21^. Injection of 2′3′-c-di-AMP also efficiently protected flies against systemic infection by DCV. However, the strongest protection was achieved following injection of 2′3′-c-di-GMP (Fig. 5A). No protection was observed in *dSTING* mutant flies, whatever the CDN injected, confirming that the four CDNs act as STING agonists. Injection of 2′3′-c-di-GMP also resulted in better protection against the rhabdovirus Vesicular Stomatitis Virus (VSV), as attested by reduced viral RNA load 4-and 5-days post-infection (Fig. 5B). Accordingly, 2′3′-c-di-GMP was a potent inducer of STING signaling, as illustrated by the strong induction of *dSTING*, *srg1*, *srg2* and *srg3* (Fig. 5C). Genome wide analysis of the transcriptional response of flies injected with 3′2′-cGAMP or 2′3′-c-di-GMP revealed that although the two CDNs upregulated the same set of genes, the response was markedly enhanced when 2′3′-c-di-GMP was used (Fig. 5D). We conclude that 2′3′-c-di-GMP is a novel potent agonist of STING in *Drosophila*, with functional relevance for the induction of antiviral immunity.

We next tested the capacity of 2′3′-c-di-GMP to trigger antiviral immunity in the three species in which we failed to see an anti-DCV effect with 2′3′-cGAMP, 2′3′-c-di-AMP or 3′2′-cGAMP (Fig. 1B). Injection of 2′3′-c-di-GMP, but not 3′2′-cGAMP, resulted in significant inhibition of DCV replication in *D. serrata, D. sechellia* and *D. mojavensis*, 2-and 3-days post-infection (Fig. 6A–C). Accordingly, genome wide analysis of the transcriptome of *D. serrata* flies injected with CDNs revealed that, while 3′2′-cGAMP did not induce large scale gene expression compared to injection of Tris buffer, 2′3′-c-di-GMP induced a robust transcriptional response (Fig. 6D).

**Figure 6:**
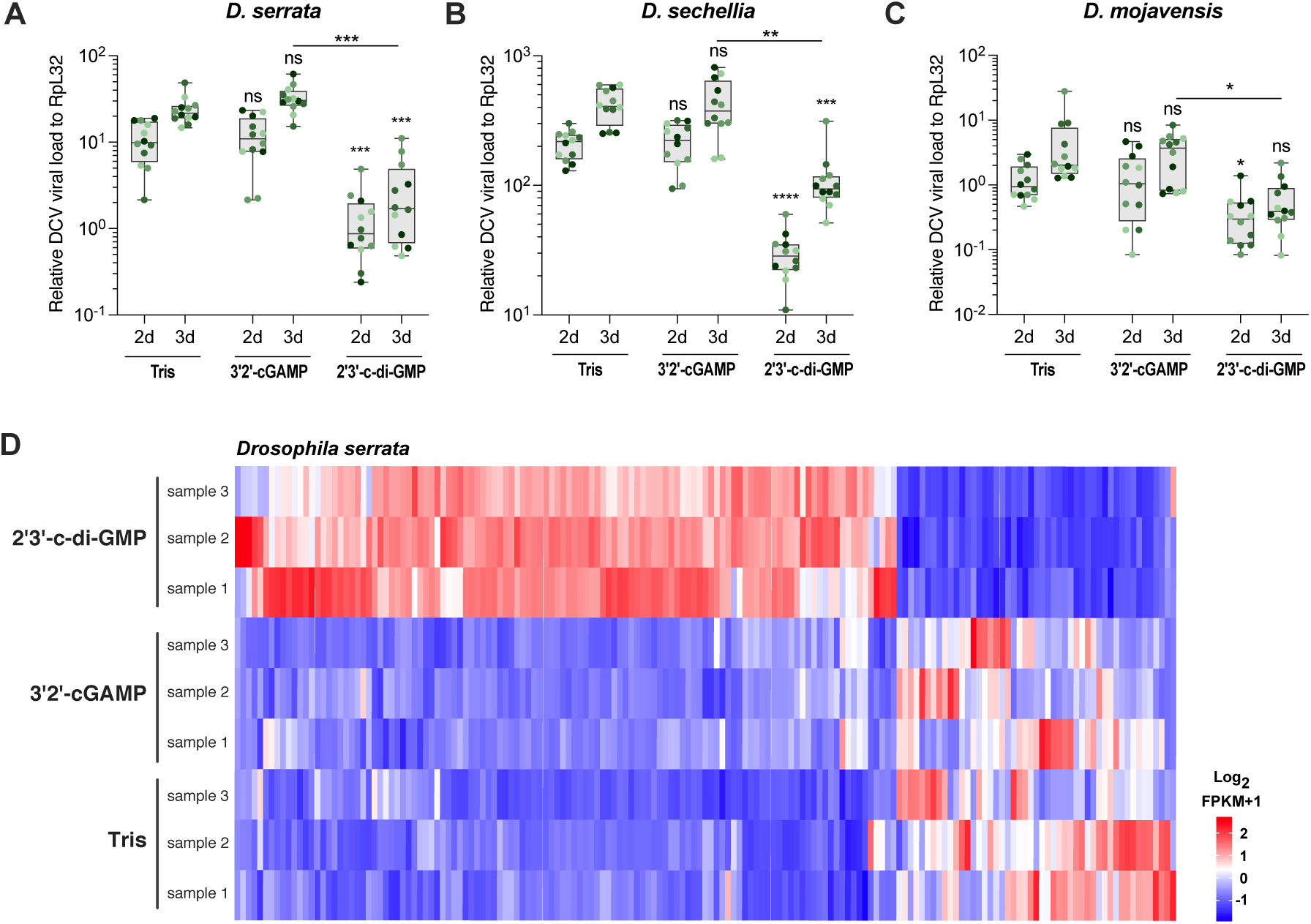
2’3’-c-di-GMP triggers antiviral immunity in *D. serrata*, *D. sechellia* and *D. mojavensis.* **A-C.** Relative DCV RNA loads at two or three days post-DCV infection in *D. serrata* (**A**), *D. sechellia* (**B**) and *D. mojavensis* (**C**) flies pre-injected with Tris or the indicated CDNs. Data are from four independent experiments. Data are shown with boxplot with scatter plot and were analysed using pairwise permutation test with FDR method (ns= non significant, *<0.05, **<0.01 and ***<0.001). **D.** Expression profiles of *D. serrata* flies 7 days after injection with Tris, 3′2′-cGAMP or 2′3′-c-di-GMP. All differentially expressed genes (DEGs) including upregulated and downregulated genes are shown in log2(FPKM+1). Data are from three independent experiments.

To understand how *Drosophila* STING recognizes 2′3′-c-di-GMP as a potent agonist, we modeled dSTING–2′3′-c-di-GMP interactions based on the dSTING–3′2′-cGAMP structure^21^. Our predictive modeling indicates that there are no steric clashes between dSTING residues and 2′3′-c-di-GMP in the CDN binding pocket and that the specific contacts which control 3′2′-cGAMP recognition also have the potential to mediate 2′3′-c-di-GMP binding (Fig. S9A). Specifically, our model predicts that each nucleobase of 2′3′-c-di-GMP stacks between Y164 and R234 residues extending from each protomer of the STING dimer and N159 coordinates the free 3′-OH of one ribose. In the dSTING–3′2′-cGAMP complex, E257 from one protomer contacts the guanosine N2 position. In our modeled dSTING–2′3′-c-di-GMP complex, this contact is conserved and E257 from the opposing protomer is predicted to additionally readout the N2 position of the second guanosine. We also observe that in our predicted dSTING–2′3′-c-di-GMP complex, T260 from one protomer coordinates the N2 position of one guanosine nucleobase, forming an additional contact that could potentially increase the specificity of dSTING for 2′3′-c-di-GMP. Residues N159, Y164, R234, E257, and T260 are highly conserved throughout the *Drosophila* genus, suggesting that 2′3′-c-di-GMP has the potential to serve as a potent agonist in most fly species (Fig. S9B). We note that 2′3′-c-di-GMP adopts a compact regiochemical conformation similar to other CDNs with mixed 2′−5′ and 3′−5′ phosphodiester linkages, including 3′2′-cGAMP and 2′3′-cGAMP, supporting the formation of a tightly closed STING dimer critical for downstream immune signaling^37, 38^. Residues controlling the specific recognition of 3′3′-linked CDNs by bacterial and animal STINGs^26, 39, 40^ are absent in *Drosophila* STING, explaining the inability of the 3′3′-c-di-GMP isomer to trigger STING antiviral immunity in flies. We finally note that our analysis of *Drosophila* STING does not suggest a structural explanation for the inability of 3′2′-cGAMP to induce antiviral immunity in certain species, as the residues controlling specific recognition of this molecule are widely conserved^21^ (Fig. S9B).

### Rapid evolution of cGLR-STING signaling in Drosophila

*Drosophila* species possess a remarkable number of cGLR genes, and we previously reported that individual species are predicted to encode as many as seven enzymes^21^. In total, we identified 207 genes encoding cGLRs in *Drosophila* species, including 61 cGLR genes in the 14 species we focused on for our *in vivo* analysis (Fig. 7A,B). These cGLRs belong to 3 major clades, with evidence of multiple gene duplication and loss events. Both *Dm*-cGLR1 and *Dm*-cGLR2 belong to the first clade in which they define two subclades. The cGLR1 subclade contains 16 genes from our species of interest, one in each of the 14 species, with a duplication in the *Drosophila* subgenus (*D. mojavensis* and *D. hydei*). The cGLR2 subclade results from a duplication of cGLR1 in the *Sophophora* subgenus and contains 10 genes from our species of interest (Fig. 1A, Fig. 7A). The major clade, with 26 genes from our species of interest, includes the *D. melanogaster* gene *CG7194*, the closest homolog of cGAS in *Drosophila* which we refer to as “cGLR3”. Members from this clade are present in all of the species we focused on in this study, except *D. virilis*, with a number of genes varying between 1 (*D. melanogaster*, *D. simulans*, *D. sechellia*, *D. mojavensis* and *D. hydei*) and 4 (*D. serrata*, *D. kikkawai*, and *D. suzukii*), and evidence for multiple duplications and losses. Finally, members of the third clade are present in 9 of our species of interest, revealing independent losses in different branches (Fig. 7A). Interestingly, enzymes from this clade exhibit a low predicted isoelectric point (pI), unlike the other cGLRs and mammalian cGAS, which have a high pI reflecting the presence of a long positively charged surface accommodating nucleic acid ligands (Suppl. Fig. 10).

**Figure 7:**
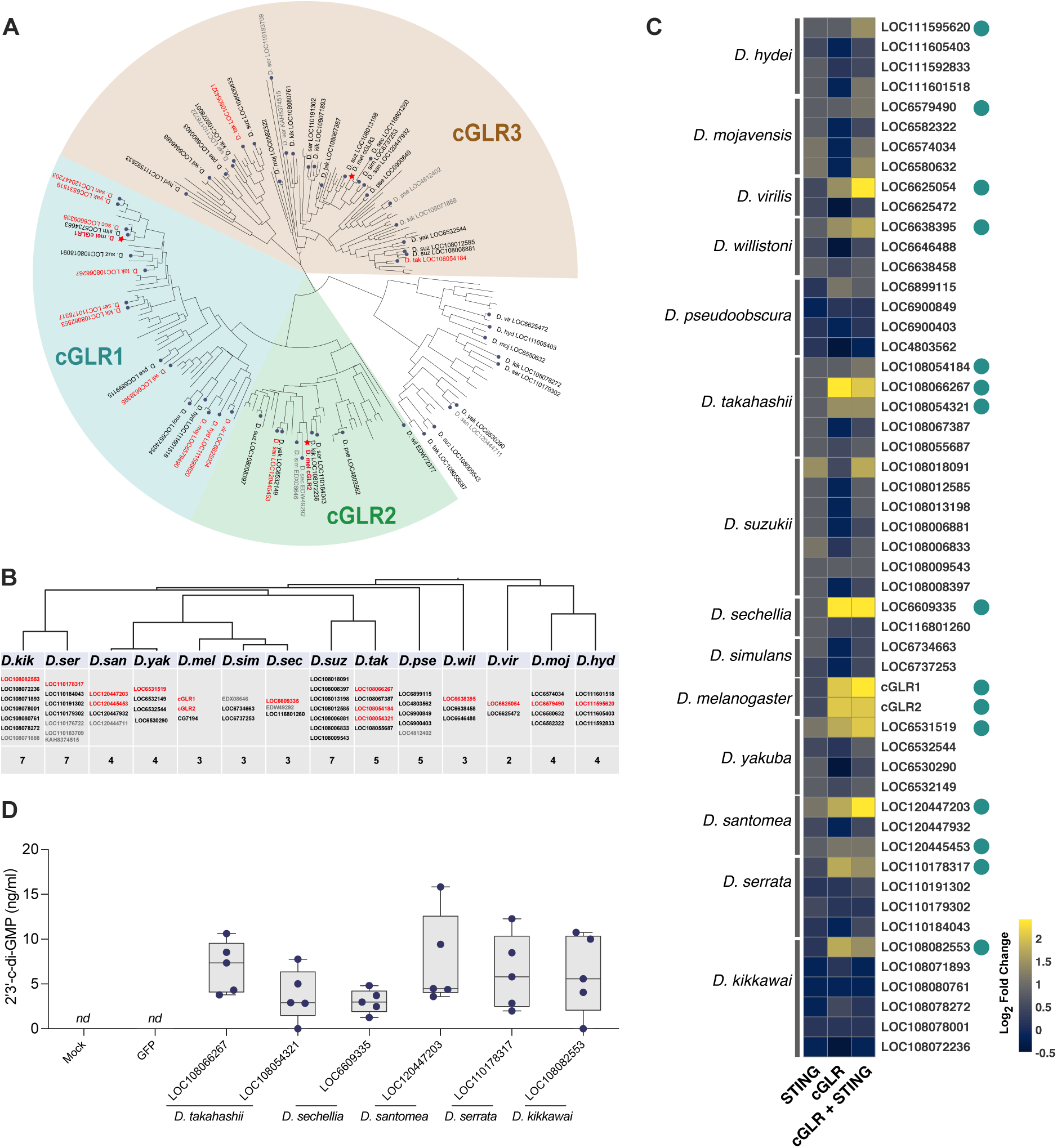
Rapid evolution of cGLR-STING signaling in *Drosophila*. **A.** Phylogenetic tree of the cGLRs identified in *Drosophila* genomes (n= 207). Sequences of predicted cGLR were aligned using MAFFT and the aligned sequences were used to construct the tree. The names of the cGLRs from the 14 *Drosophila* species are indicated and those able to activate STING signaling are labelled in red while the inactive ones are in black. The red stars indicate cGLRs from *D. melanogaster*. The 8 cGLRs in grey were not tested. **B.** The predicted cGLRs encoded in the genome of the 14 *Drosophila* species. The cGLRs able to activate STING signaling are labelled in red. **C.** Heatmap showing log2 fold change intensities in *dSTING* luciferase reporter gene activity in S2 cells expressing the indicated 52 candidate cGLRs in comparison to control. Four conditions were tested with empty vector, STING, cGLR and cGLR+STING from the same species. 15 positive candidates highlighted with dots were selected by mixed-effect model (RELM) with *p* value ≤0.05 (Tukey correction for multiple hypothesis testing) and with fold change > 2 with cGLR or cGLR with STING conditions. Data are from 2 to 4 independent experiments. **D.** Concentration of 2′3′-c-di-GMP measured by LC-MS/MS in lysates from *dSTING* knockout S2 cells ectopically expressing the indicated cGLRs. Data are from 5 independent experiments.

Expression vectors for 52 cGLR genes from the 14 species we focused on in this study were constructed and transfected in S2 cells together with vectors expressing STING from the corresponding species and a *dSTING-luciferase* reporter plasmid, to monitor activation of the pathway^15^. We identified 15 cGLRs, including *D. melanogaster* cGLR1 and cGLR2, that are able to activate STING signaling in S2 cells (Fig. 7C). Most of them (13/15) belong to the cGLR1/2 clade. The two other active cGLRs, both from the species *D. takahashi*, belong to the same clade as the *D. melanogaster* gene *CG7194* (*cGLR3*) (Fig. 7A). Of note, several of them, including the cGLR1 orthologs from *D. sechellia* and *D. serrata*, produce 2′3′-c-di-GMP when transfected into S2 cells (Fig. 7D). Overall, our findings confirm that cGLRs from different clades share the property to activate STING signaling and point to a dynamic evolution of these receptors in the *Drosophila* lineage, with 2′3′-c-di-GMP playing a major role at least in some species.

## Discussion

We show here that CDNs are produced in *D. melanogaster* flies in response to virus infection, thus connecting sensing of viral infection to the enzymatic activity of cGLRs *in vivo*. Importantly, we show that a set of CDNs, rather than a single molecular species, are produced during infection. Of these molecules, the most potent STING agonist is a previously unknown CDN, 2′3′-c-di-GMP. This may explain our previous observation that transgenic flies ectopically expressing cGLR2, which produce this CDN, are better protected against DCV infection than flies ectopically expressing cGLR1, which do not produce 2′3′-c-di-GMP^20^. Our results contrast with the situation in mammals, where cGAS primarily synthesize 2′3′-cGAMP^29, 32^, and highlights intriguing similarities with the CBASS system in bacteria, where CD-NTase enzymes can produce a broad diversity of cyclic di-or tri-nucleotides^23, 25, 41, 42^. This diversity has been proposed to give an edge to bacteria to escape viral nucleases targeting them. Indeed, 3′2′-c-GAMP is resistant to both poxin and Acb1 nucleases, which are produced respectively by insect DNA viruses and phages and readily cleave canonical 2′–5′ linkage in CDNs^43–46^. An alternative explanation for the diversity of CDNs produced in *Drosophila* may be the existence of additional sensors, besides STING, which may preferentially recognize some species of CDNs. Indeed, the only known CDN receptor in mammals, besides STING, is RECON, which preferentially binds 3′– 5′-linked CDNs and CTNs, rather than 2′3′-cGAMP^23, 47^. However, these alternative sensors for CDNs remain unknown in *Drosophila*, and we note that the antiviral protection induced by the four CDNs we identified is fully dependent on STING. A third possibility could be related to extracellular functions of CDNs in flies. Indeed, we and others had previously shown that injection of CDNs into the fly body cavity results in potent STING-dependent antiviral protection, pointing to the existence of import mechanisms for CDNs^16, 19, 21^. We now show that CDNs are found circulating in the hemolymph upon viral infection, suggesting that CDN export mechanisms may operate in virus-infected cells. This would provide an efficient means to trigger a systemic response to infection and one may hypothesize that some CDNs are more efficiently transported in and out of cells and function as immuno-transmitters, whereas others function as *bona fide* second messengers in the cells in which they are produced. The observation that 3′3′-c-di-GMP protects *D. melanogaster* against enteric infection when fed to flies^16^ but has no effect on systemic antiviral immunity when it is injected in the body cavity^16, 19^, supports the hypothesis that regulation of the transport across cellular membranes plays a role in the biology of CDNs. Finally, an additional possibility is that the diversity of CDNs produced reflects the number of cGLRs that produce them, possibly in different tissues.

The genes encoding cGLRs in *Drosophila* have experienced multiple duplication and losses, resulting in a number of receptors ranging from 2 to 7 according to the species and defining three major clades. Some 30% of the cGLRs tested (15 out of 52) can produce STING agonists in *Drosophila* S2 cells and most of the active enzymes we identified belong to the clade encompassing *D. melanogaster* cGLR1 and 2, with only two active enzymes identified in the clade encompassing the third cGLR in *D. melanogaster*, cGLR3. Our data reveal that the activity of *D. pseudoananassae* cGLR2, like that of *D. melanogaster* cGLR1, can be activated *in vitro* by dsRNA, confirming that cGLRs function as PRRs and that the clade to which cGLR1 and cGLR2 belong is primarily dedicated to sensing of viral RNAs. The observation that significant amounts of CDNs are still induced in response to DCV infection in *cGLR1* or *cGLR2 D. melanogaster* mutant flies, confirm that both receptors can be independently activated by the virus and contribute to the control of the infection.

Intriguingly, we noted residual production of CDNs in DCV-infected *cGLR1/2* double mutant flies, which suggests that cGLR3 (CG7194) may also be activated upon sensing viral infection in *D. melanogaster*. This hypothesis is supported by the observation that two members of the cGLR3 clade in *D. takahashi* can activate STING in transfected cells and that one of them produces 2′3′-c-di-GMP.

cGLRs within the cGLR1/2 clade and the cGLR3 clade are defined by characteristic positive residues in the ligand binding surface that control the recognition of negatively charged nucleic acids^20, 21^. These residues contribute to an overall basic pI, consistent with other nucleic acid sensing cGLRs including cGAS^40^. Intriguingly, we find that a third clade of *Drosophila* cGLRs have calculated pIs <6 and are predicted to have neutral and acidic ligand binding surfaces, suggesting they may not recognize nucleic acid ligands. No representative members of this clade have been characterized, and we predict these proteins could function as PRRs to detect non-nucleic acid pathogen associated molecules. Whether the lack of activity of the other cGLRs we tested here reflects the absence of activating ligands in S2 cells or the fact that their products cannot bind to and activate STING remains an open question, the answer to which will provide important insight on the biology of cGLRs. Interestingly, similar species-specific differences between the activity of cGAS from different mammals, including within the primate lineage, have recently been described and shown to result, at least in part, from differences of subcellular localization of the receptor^48^.

Our findings nicely complement a recent study that identified more than 3,000 cGLRs in the genomes of most metazoans and characterized a subset of them as PRRs responding to dsRNA or dsDNA^40^. While most metazoan cGLR synthesize 2′3′-cGAMP, some cGLRs also produce alternative nucleotide signals including isoforms of c-UMP–AMP with pyrimidine bases, and it will be interesting to see if some the *Drosophila* cGLRs can also synthesize still additional CDNs besides the four we identified here. Overall, our results create a new foundation to better understanding of the biology of cGLRs using the *Drosophila* model. Further experiments using *cGLR1/2* double mutant flies or even *cGLR1/2/3* triple mutant flies challenged by a panel of biotic or abiotic stresses may provide some clue regarding their function in a physiological setting. In addition, genetic complementation of cGLR mutant *D. melanogaster* with genes encoding cGLRs from other species is expected to shed light on their function, with possible implications for the function of uncharacterized cGLRs with complete catalytic sites encoded in human and other vertebrates (e.g., MB21D2)^21^.

## Acknowledgements

We thank Joao Marques for critical reading of the manuscript, Hunter Toyoda for help purifying proteins, and members of the Kranzusch, Imler and Cai labs for discussions and helpful comments. JLI acknowledges financial support from CNRS and grants from ANR (ANR-10-IDEX-0002, ANR-11-EQPX-0022, ANR-17-EURE-0023, ANR-20-SFRI0012, ANR-22-CE15-0019) HC was supported by the Natural Science Foundation (32000662), Guangdong Provincial Science Fund for Distinguished Young Scholars (2023B1515020098), Guangdong Provincial Young Scholars academic exchange program (2022A0505030018), Youth Talent Support Programme of Guangdong Provincial Association for Science and Technology (SKXRC202229), Science and Technology Program of Guangzhou (202102020090). DC was supported by Natural Science Foundation of China (NSFC, 32200578). RJ was supported by the National Key R&D Program of China (2021YFA0805800, 2020YFA0803202), the National Natural Science Foundation of China (31970538) and the Local Innovative and Research Teams Project of Guangdong Perl River Talents Program (2017BT01S155). JLI, DC and RJ. Acknowledge support by the Chinese National Overseas Expertise Introduction Center for Discipline Innovation (Project ‘111’ (D18010). PJK acknowledges financial support from the Pew Biomedical Scholars program, the Burroughs Wellcome Fund PATH program, The Mathers Foundation, The Mark Foundation for Cancer Research, the Cancer Research Institute, the Parker Institute for Cancer Immunotherapy, and the National Institutes of Health (1DP2GM146250-01). KMS is supported as an NCI F99 Graduate Fellow NIH 1F99CA274660-01.

## STAR METHODS

### Cell lines

HEK293T (ATCC) were cultured in Dulbecco’s modified Eagle medium (DMEM) (Sigma-Aldrich) supplemented with 10% fetal bovine serum (FBS) (Excell Bio), 100 U mL^−1^ penicillin (Sigma-Aldrich) and 100 µg mL^−1^ streptomycin (Sigma-Aldrich) at 37°C and 5% CO_2_. Schneider 2 (S2) cells (Invivogen) were cultured in Schneider’s *Drosophila* Medium (Thermal Fisher Scientific) supplemented with 10% FBS, 100 U mL^−1^ penicillin (Sigma-Aldrich) and 100 µg mL^−1^ streptomycin (Sigma-Aldrich) at 27°C. To generate Sting knockout cells by CRISPR-Cas9, three CRISPR RNAs targeting the first translated exon in the *D. melanogaster Sting* gene were cloned into the pAc-sgRNA-Casp vector^1^. After transfection, S2 cells were grown under selection with 5 µg mL^−1^ puromycin (Invitrogen) for two weeks. Knockout of Sting was verified by Sanger sequencing.

### Drosophila strains

Fly stocks were raised on standard cornmeal agar medium at 25°C and were free of *Wolbachia*. All fly stocks used (*DrosDel w*^1118^, *dSTING^L76GfsTer^*^11^, *dSTING^Rxn^*, *dSTING^Control^*, cGLR1 or cGLR2 single knockout or double knockout flies, the transgenic lines of wild-type and AFA mutant versions of cGLR1 and cGLR2, and the GFP control) have been described previously^1–3^. The different species of *Drosophila* were kindly provided by Prof. Jian Lu, Peking University (*D.willistoni*, *D.yakuba*, *D.simulans*, *D.virilis* and *D.takahashi*); Prof. Binyan Lu, Chinese Academy of Sciences (*D.suzukii* and *D. hydei*); Prof. Shuoyang Wen, South China Agricultural University (*D.serrata* and *D.kikkawai*); Prof. Wei Wu, Chinese Academy of Sciences (*D.pseudoobscura*); Prof. Yufeng Pan, Southeast University (*D.mojavensis*) and Prof. Nicolas Gompe, Lüdwig Maximilian University (*D.sechelia* and *D.santomea*).

### Plasmids

Coding sequences of predicted cGLRs were PCR-amplified from the cDNA prepared from flies from the different *Drosophila* species using 2×EasyTaq PCR SuperMix (TransGen Biotech, China) and cloned into the pAc5.1 vector using ClonExpress MultiS (Vazyme, China). The coding sequences of STING from fourteen species of *Drosophila* were synthesized by the IGE biotechnology company (China) and sub-cloned to pAc5.1 vector.

### Antiviral effect of cyclic di-nucleotides in fourteen species of Drosophila

Cyclic dinucleotides including 3′2′-cGAMP (Biolog C238), 2′3′-cGAMP (Biolog C161), 2′3′-c-di-AMP (Biolog C187), 2′3′-c-di-GMP (Biolog C182) or 3′3′-c-di-GMP (Biolog C057) were dissolved in 10 mM Tris-HCl pH 7.5 and diluted to the indicated concentrations. 3–5 days old male flies were injected with 69 nL of cyclic dinucleotide solution or 10 mM Tris-HCl pH 7.5 (negative control) by intrathoracic injection using a Nanoject II apparatus (Drummond Scientific). 7 days post CDN injection, the flies were injected with 4.6 nL of DCV (50 PFU for all the tested species except *D. melanogaster* and *D. yakuba*, for which the dose was 5 PFU) in 10 mM Tris-HCl pH 7.5. Flies were collected 48 h and 72 h later in pools of 6 males and homogenized for RNA extraction and RT-qPCR analysis, as described.

### Fly hemolymph and lysate preparation for Liquid Chromatography Mass Spectrometry (LC-MS)

For hemolymph collection, anesthesized adult flies were punctured using a tungsten needle and transferred to a 0.5 mL microcentrifuge tube (12 flies/tube) with a hole pierced in the bottom. The tube with flies was then transferred to a 1.5 mL Eppendorf tube. The two-tube assembly was then centrifuged for 10 min at 2,300g at 4°C. The procedure was repeated until the hemolymph of one thousand flies was collected. Hemolymph was stored at −80°C.

For lysate preparation, 500 adult flies were collected in five 2 mL microfuge tubes (100 flies/ tube). 10 small zirconia beads were added per tube together with 100 μL precooled extraction reagent (2/2/1 [v/v/v] methanol, acetonitrile, water mixture) with 10 ng mL^−1^ 3’3’-cAIMP as an internal standard in the LC-MS/MS analysis. The tubes with flies were frozen in liquid nitrogen and then homogenized using Precellys Evolution homogenizer (4°C, 5800rpm, 2*30s pause 12s). The tubes were then centrifuged for 10 min at 12,000g at 4°C. The supernatant was collected into a new microfuge tube, which was centrifuged again. The collected supernatant was transferred into 2 mL safe-lock tubes and heated at 95°C for 10 min, before cooling on ice. 500 µL chloroform was then added to the tubes, which were vortexed and placed at −20°C for 10 min. Subsequently, the tubes were centrifuged at 20,000 x g for 15 min, and the supernatants was transferred into fresh microfuge tubes. Next, the lysate was loaded on HLB SPE columns. The eluents were collected and then concentrated by evaporation and resuspended in 200 μL 0.1% formic acid water for LC-MS/MS analyses.

### Identification of CDN using LC-MS

To detect the CDNs production in the lysate of DCV infected flies, high resolution LC-MS analysis was performed using a Ultimate3000 (ThermoFisher Scientific) coupled to a Quadrupole-Orbitrap Hybrid mass spectrometer (Q-Exactive, ThermoFisher Scientific). A volume of 5μL sample was injected into a Agilent SB-Aq RRHD column (1.8μm 2.1*100mm) carried out at 40°C. The mobile phase consisted of 5 mM ammonium carbonate (A) and acetonitrile (B). The HPLC gradient: 0-14% B in 5.0 min, 14-25% B in 7 min, 25–100% B in 7.1 min, 100% B in 10.8 min, 100-0% B in 11.0 min, 0% B in 14.0 min; 0.300 mL min^−1^. Full scan MS spectra were acquired in a mass range from m/z 300 to 1100 with a resolution of 70000 and a AGC target of 1.00E+06 in the Orbitrap mass analyser. The target ions were sequentially isolated for HCD fragmentation and progeny ions were detected in orbitrap with PRM mode. The parent ion was isolated with an isolation window of 0.4 m/z units, fragmented (Resolution = 17500, nce=20, Maximum Inject Time: 50ms). Identification of CDNs was performed by targeted mass analysis for exact masses and formulae for all possible CDNs.

### Measurement of CDN in the hemolymph or lysate

For measurement of 2’3’-cGAMP, 3’2’-cGAMP, 2’3’-c-di-AMP and 2’3’-c-di-GMP in the hemolymph, samples were analyzed using an Agilent 1290 System coupled to an Agilent 6470. A volume of 5μL was injected into a Nucleodur Pyramid C18 column (50 x 3 mm; 3 μm Macherey Nagel, Duren, Germany) carried out at 40°C. The mobile phase consisted of 0.1% formic acid in acetonitrile (A) and 0.1% formic acid in water (B). HPLC gradient are described in Supplementary Table 1. The analytes were ionized by means of electro spray ionization in positive mode with the Delta EMV at 820 V. The source parameters were described in Supplementary Table 2. For each CDN, the MRM transition(s) (m/z), Dwell, Frag (V) and CE (V) Cell acc (V) were described in Supplementary Table 3.

For measurement of 2’3’-cGAMP, 3’2’-cGAMP, 2’3’-c-di-AMP, 2’3’-c-di-GMP in whole fly lysates, the samples were analyzed using aAgilent 1290 System coupled to a Agilent 6470. A volume of 5 μL was injected into a Nucleodur Pyramid C18 column (50 x 3 mm; 3 μm Macherey Nagel, Duren, Germany) carried out at 40°C. The mobile phase consisted of 0.1% formic acid in acetonitrile (A) and 0.1% formic acid in water (B). The HPLC gradient and flow rate were described in Supplementary Table 4. The analytes were ionized by means of electro spray ionization in positive mode with the Delta EMV at 820 V. The source parameters were described in Supplementary Table 5. For each CDN, the MRM transition(s) (m/z), Dwell, Frag (V) and CE (V) Cell acc (V) were described in Supplementary Table 6.

### D. melanogaster cyclic dinucleotide injection and signaling analysis

Cyclic dinucleotides including 2’3’-c-di-GMP (Biolog C182), 2’3’-c-di-AMP (Biolog C187), 3′2′-cGAMP (Biolog C238), 2′3′-cGAMP (Biolog C161) and 3′3′-c-di-GMP (Biolog C057) were dissolved in 10 mM Tris-HCl pH 7.5 and diluted to the indicated concentrations. 3–5-day old adult flies were injected with 69 nL of cyclic dinucleotide solution or 10 mM Tris-HCl pH 7.5 (negative control) by intrathoracic injection using a Nanoject II apparatus (Drummond Scientific). Flies were collected 7 days later in pools of 6 individuals (3 males and 3 females) and homogenized for RNA extraction and quantitative PCR with reverse transcription (RT–qPCR) analysis, as described.

### Bioinformatics and Drosophila cGLR and STING sequence analysis

Building on previous analyses^4–7^, *Drosophila* cGLRs were identified using the amino acid sequence of *D. melanogaster* cGLR1 (NP_788360.2) to seed a position-specific iterative BLAST (PSI-BLAST) search of *Drosophila* genomes (taxid:7215) in the NCBI non-redundant protein database. The PSI-BLAST search was performed with an E value cutoff 0.005 for inclusion into the next search round, BLOSUM62 scoring matrix, gap costs settings existence 11 and extension 1, and using conditional compositional score matrix adjustment. Iterative PSI-BLAST search was performed for 5 rounds and candidate *Drosophila* cGLR sequences were collected. Putative cGLR protein sequences were aligned using MAFFT (FFT-NS-i iterative refinement method)^8^; this alignment was used to construct a phylogenetic tree in Geneious Prime v2022.0.1 using the neighbor-joining method and Jukes-Cantor genetic distance model with no outgroup. Candidate proteins were analyzed by clade and selected for known cGLR domain organization and predicted structural homology to *T. castaneum* cGLR (PBD: 7LT2), including the presence of a conserved nucleotidyltransferase domain with a G[S/G] activation loop and [E/D]h[E/D] _X50–90_ [E/D] catalytic triad^9^. Manual analysis and curation of candidate cGLR sequences was performed based on alignments and predictive structural homology using Phyre2^10^ and AlphaFold^11^. Manual refinement was also used to exclude duplicate sequences, gene isoforms, and proteins less than 250 residues. All cGLR sequences in the final tree were accessed from NCBI March 26^th^, 2023. The *D. bipectinata* cGLR2 sequence shown in Supplementary Figure 7A,B and used for all biochemistry experiments was accessed from NCBI in January, 2020 under accession code XP_017096409.1. NCBI available genomes from 49 species in the *Drosophila* genus are represented in the final tree. iTOL was used for tree visualization and annotation^12^, including annotation of all cGLRs identified in fourteen *Drosophila* species of interest to this study. Isoelectric point was predicted by Geneious Prime software. Clustering of sequences in the final unrooted tree was used to define clades related to *D. melanogaster* cGLR1 (NP_788360.2), cGLR2 (A8DYP7.2), and cGLR3 (CG7194; AAF50449.1). The cGLR2-related clade was identified by the presence of *D. melanogaster* cGLR2 (A8DYP7.2) and extracted from the full cGLR tree for alignment using MAFFT (FFT-NS-i iterative refinement method), shown in Supplementary Figure 7A. cGLR2 sequences of interest in Supplementary Figure 7B were separately aligned and the secondary structure of *D. pseudoananassae* cGLR2 was generated based on the AlphaFold predicted structure.

*Drosophila* STING proteins were identified as a subset of eukaryotic STING proteins identified in ref. ^6^. STING sequences were accessed from NCBI May 2022. *Drosophila* STING protein sequences from fourteen species of interest were aligned using MAFFT (FFT-NS-i iterative refinement method)^8^. Alignment in Supplementary Figure 9B is shown in relation to the secondary structure of the *D. eugracilis−*3′2′-cGAMP complex (PDB: 7MWZ)^5^.

### Screening of cGLRs based on cellular STING signaling assays

To test the activation of STING promoter luciferase reporter by predicted cGLRs, 96-well tissue culture plates were seeded with 2.5 x 10^5^ S2 cells per well. After 3h, each well was transfected with 200ng pGL3 plasmid expressing firefly luciferase under transcriptional control of the Sting promoter, 25ng pAc5.1 plasmid constitutively expressing Renilla luciferase, 75ng pAc5.1 plasmid expressing predicted cGLRs, 25ng pAc5.1 plasmid expressing Sting, finally empty Ac5.1 plasmid to reach a total amount of 325ng plasmid for each two wells. All transfections of S2 cells were performed using lipofectamine 3000 (Invitrogen, Thermo Fisher Scientific) according to the manufacturer’s instructions. After 48 hours of transfection, cells were lysed in 100µL Lysis Buffer (Promega) per well. Firefly and Renilla luciferase activity was measured on 75µL lysate using the Dual-Luciferase® Reporter Assay System (Promega).

### RNA-Sequencing of D. melanogaster injected with CDNs

Male flies of *D. melanogaster* or *D. serrata* were injected with 69 nL/fly of either 10 mM Tris (pH 7.5), 2’3’-cGAMP (0.9 mg mL^−1^) or 2’3’-cdi-GMP (0.9 mg mL^−1^) by intrathoracic injection (Nanoject II apparatus) in three independent experiments. Injected flies were collected in pools of 6 individuals at 7 days post injection. Total RNA was isolated from injected flies using TRIzol™ Reagent (Invitrogen), according to the manufacturer’s protocol. RNA quality was assessed on an Agilent 2100 Bioanalyzer (Agilent Technologies, Palo Alto, CA, USA) and checked using RNase free agarose gel electrophoresis. After total RNA was extracted, eukaryotic mRNA was enriched by Oligo(dT) beads. Then the enriched mRNA was fragmented into short fragments using fragmentation buffer and reverse transcribed into cDNA by using NEBNext Ultra RNA Library Prep Kit for Illumina (NEB #7530, New England Biolabs, Ipswich, MA, USA). The purified double-stranded cDNA fragments were end repaired, A base added, and ligated to Illumina sequencing adapters. The ligation reaction was purified with the AMPure XP Beads (1.0X). Ligated fragments were subjected to size selection by agarose gel electrophoresis and polymerase chain reaction (PCR) amplified. The resulting cDNA library was sequenced using Illumina Novaseq6000 by Gene Denovo Biotechnology Co. (Guangzhou, China).

### Transcriptome analysis

After filtering by fastp^13^ (version 0.18.0), reads were mapped using HISAT2. 2.4^14^ with “-rna-strandness RF” and other parameters set as a default to the genome of *D. melanogaster* (Ensembl_release102) and *D. serrata* (GCF_002093755.1). RNAs differential expression analysis was performed by DESeq2^15^ software between two different groups (and by edgeR between two samples). The genes/transcripts with the parameter of false discovery rate (FDR) below 0.05 and absolute fold change≥2 were considered differentially expressed genes/transcripts.

### Protein expression and purification

Recombinant cGLR proteins were expressed and purified using methods previously optimized for human cGAS and *Drosophila* cGLRs, as described previously^5, 16^. Full length cGLR coding sequences were codon-optimized for expression in *E. coli* and cloned from synthetic constructs (GeneArt or Integrated DNA Technologies) into a custom pET16 expression vector with an N-terminal 6×His-MBP fusion tag. Briefly, transformed BL21-CodonPlus (DE3)-RIL *E. coli* (Agilent) were grown in MDG media overnight prior to inoculation of M9ZB media at an OD_600_ of 0.0475. M9ZB cultures were grown to OD_600_ of 2.5 (approximately 5 h at 37°C with shaking at 230 rpm) followed by cooling on ice for 20 min. Cultures were induced with 500 μM IPTG prior to incubation at 16°C overnight with shaking at 230 rpm. Cultures were pelleted the following day and either flash frozen in liquid nitrogen for storage at −80°C or directly lysed for purification.

For large-scale protein purification, proteins were expressed with a 6×His-MBP fusion tag and grown as ∼4–8× 1 L cultures in M9ZB media. Pellets were lysed by sonication in lysis buffer (20 mM HEPES pH 7.5, 400 mM NaCl, 30 mM imidazole, 10% glycerol and 1 mM DTT) and clarified by centrifugation at ∼47,850 × g for 30 min at 4°C and subsequent filtration through glass wool. Recombinant protein was purified by gravity-flow over NiNTA resin (Qiagen). Resin was washed with lysis buffer supplemented to 1 M NaCl and then eluted with 20 mL of lysis buffer supplemented to 300 mM imidazole. MBP-tagged fusion proteins were buffer exchanged into lysis buffer with 4% glycerol and no imidazole to optimize conditions for overnight cleavage by recombinant TEV protease at ∼10°C. cGLR proteins were next purified by ion exchange chromatography using 5 mL HiTrap Heparin HP columns (GE Healthcare) and eluted across a 150–1000 mM NaCl gradient in buffer with 10% glycerol. Target protein fractions were pooled concentrated to ∼10–30 mg mL^−1^ and flash-frozen with liquid nitrogen and stored at −80°C for biochemistry experiments.

### Nucleotide product synthesis analysis

cGLR nucleotide synthesis activity was analyzed by thin-layer chromatography as previously described^4, 5^. For all biochemistry reactions analyzed by TLC, 5 μM recombinant protein preparations were incubated in 5 or 10 μL reactions containing 0.5 μL α-^32^P labeled ATP or GTP, 200 μM unlabeled NTPs, and 1 mM MnCl_2_ in a final reaction buffer of 50 mM Tris-HCl pH 7.5, ∼50 mM KCl (final KCl = 100 mM), 1 mM TCEP. Reactions were additionally supplemented with ∼1 μg poly I:C or 5 μM nucleic acid ligands, as indicated. Besides poly I:C all nucleic acids used in this study are 40 nucleotide (nt) or base pairs (bp) in length. Reactions were incubated at 37°C for two hours and subsequently treated with 1 μL Quick CIP phosphatase (New England Biolabs) for 20 min at 37°C to remove unreacted phosphate signal. 0.5 μL of each reaction was spotted on a 20-cm × 20-cm PEI-cellulose thin-layer chromatography plate. Plates were run with 1.5 M KH_2_PO_4_ solvent until∼2.5 cm from top of the plate, dried at room-temperature, and exposed to a phosphor-screen prior to signal detection with a Typhoon Trio Variable Mode Imager System (GE Healthcare). TLC images were adjusted for contrast using FIJI^40^ and quantified using ImageQuant (8.2.0). GTP conversion to nucleotide product formation was measured according to the ratio of product to total signal for each reaction.

Nuclease P1 cleavage analysis was performed using cGLR2 reactions labeled with either α-^32^P-ATP or α-^32^P-GTP as previously described^4, 5^. Briefly, radiolabeled nucleotide products were incubated with Nuclease P1 (80 mU, Sigma N8630) in buffer (30 mM NaOAc pH 5.3, 5 mM ZnSO_4_, 50 mM NaCl) for 30 min in the presence of Quick CIP (NEB).

### Nucleotide purification and HPLC analysis

Enzymatic synthesis of cGLR nucleotide products for HPLC analysis was performed using 100 μL reactions containing 5 μM cGLR enzyme, 100 μM ATP, 100 μM GTP, 10 μg poly I:C, 1 mM MnCl_2_, and 50 mM Tris-HCl pH 7.5. KCl was adjusted to a final concentration of 100 mM. Reactions were incubated at 37°C for 2 h and then terminated by 2 min incubation at 95°C. Nucleotide product was recovered by filtering reactions through a 30-kDa cutoff concentrator (Amicon) to remove protein. Nucleotide products were separated on an Agilent 1200 Infinity Series LC system using a C18 column (Zorbax Bonus-RP 4.6 × 150 mm, 3.5 μm) at 40°C. Products were eluted at a flow rate of 1 mL min^−1^ with a buffer of 50 mM NaH_2_PO_4_ pH 6.8 supplemented with 3% acetonitrile.

### Synthetic cyclic dinucleotide standards

Synthetic nucleotide standards used for HPLC analysis and mass-spectrometry analysis were purchased from Biolog Life Science Institute: 2′3′-cGAMP (cat no. C 161), 3′2′-cGAMP (cat no. C 238), 2′3′-c-di-AMP (cat no. C 187) and 2′3′-c-di-GMP (cat no. C 182).

### dSTING and 2′3′-c-di-GMP structural modeling

Coordinate and cif restraint files for 2′3′-c-di-GMP were generated by the eLBOW program on PHENIX^17^ using the ChemDraw v.20.0.38 generated SMILES for this ligand. The resulting files were used to model the 2′3′-c-d-GMP ligand in complex with *D. eugracilis* STING by fitting to the 3′2′-cGAMP ligand density in the dSTING*−*3′2′-cGAMP complex density map (PDB 7MWZ) using Coot^18^.

### LC-MS/MS analysis of in vitro cGLR2 reaction products

cGLR2 reaction products were generated in a 100 μL reaction with 5 μM cGLR enzyme, 200 μM GTP, 10 μg poly I:C, 1 mM MnCl_2_, and 50 mM Tris-HCl pH 7.5 and were analyzed by the commercial company MS-Omics using LC-MS/MS, as previously described (Li et al., 2023). Briefly, analysis was carried out using a Vanquish™ Horizon UHPLC System coupled to Orbitrap Exploris 240 Mass Spectrometer (Thermo Fisher Scientific, US).

First, UHPLC was performed using an Infinity Lab PoroShell 120 HILIC-Z PEEK lined column with the dimension of 2.1 × 150mm and particle size of 2.7 µm (Agilent Technologies). Mobile phase A was composed of 10 mM ammonium acetate, pH 9 in 90% Acetonitrile LC-MS grade (VWR Chemicals, Leuven) and 10% Ultra-pure water from Direct-Q® 3 UV Water Purification System with LC-Pak® Polisher (Merck KGaA, Darmstadt). Mobile phase B was composed of 10 mM ammonium acetate, pH 9 in ultra-pure water with 5 µM medronic acid (InfinityLab Deactivator additive, Agilent Technologies). The UHPLC column temperature was set at 30 °C and samples were analyzed at an injection volume of 5 µl. UHPLC was run using a flow rate kept at 250 µl mL^−1^ consisting of a 2 min hold at 10% B, increased to 40% B at 14 min, held till 15 min, decreased to 10% B at 16 min and held for 8 min.

For MS analysis, a heated electrospray ionization interface was used as ionization source and the analysis was performed in positive ionization mode from m/z 300 to 1500 at a mass resolution of 120000. Ion source parameters used: Sheath gas flow rate, 20 (arbitrary units); auxiliary gas flow rate, 5 (arbitrary units); Sweep gas flow rate, 1 (arbitrary units), capillary temperature, 350°C; S-lens radiofrequency level 70; automatic gain control (AGC) target, 1E6 (Standard); maximum injection time, 100 ms; spray voltage 3.5 kV in positive. MS2 spectra was acquired using data dependent acquisition (DDA) with the following parameters: mass resolution 45000, isolation window m/z 0.4 and normalized collision energy 20, 40 and 60 eV. Freestyle 1.4 (Thermo Fisher Scientific) was used to analyze data and generate MS/MS spectra.

### Statistical analysis

All statistical analyses were performed in R (version 4.1.2) and GraphPad Prism9. To perform the qPCR analysis, after calculating the 2ΔCt for each sample were tested by permutation test using coin package. In addition, survival analyses using Log-rank tests were performed to compare each condition to control lines. cGLRs from different species were analysed using mixed-effect model (RELM) with Tukey correction of the log transformed data. Fold change is determined in comparison to the transfected control (pAc5.1). 15 cGLRs from different species were selected with *p* value :: 0.05 with fold change ý 2.

---

**Table S1.** HPLC gradient.

|  | **Time** | **Mobile phase (A)** | **Mobile phase (B)** | **Flow** |
| --- | --- | --- | --- | --- |
| **1** | 0.00min | 4.00% | 96.00% | 0.15 mL/min |
| **1** | 5.00 min | 4.00% | 96.00% | 0.20 mL/min |
| **2** | 5.10 min | 95.00% | 5.00% | 0.50 mL/min |
| **3** | 8.50 min | 95.00% | 5.00% | 0.50 mL/min |
| **4** | 8.70 min | 10.00% | 90.00% | 0.40 mL/min |
| **5** | 13.00 min | 10.00% | 90.00% | 0.40 mL/min |
| **6** | 13.20 min | 4.00% | 96.00% | 0.15 mL/min |
| **7** | 13.80 min | 4.00% | 96.00% | 0.15 mL/min |

**Table S2.** The source parameters.

| **Parameter** | **Value (+)** | **Value (-)** |
| --- | --- | --- |
| **Gas temp (°C)** | 350 | 350 |
| **Gas Flow (L min<sup>-1</sup>)** | 7 | 7 |
| **Nebulizer (psi)** | 35 | 35 |
| **SheathGasHeater** | 350 | 350 |
| **SheathGasFlow** | 12 | 12 |
| **Capillary (V)** | 2500 | 3500 |
| **VCharging** | 800 | 500 |

**Table S3.**
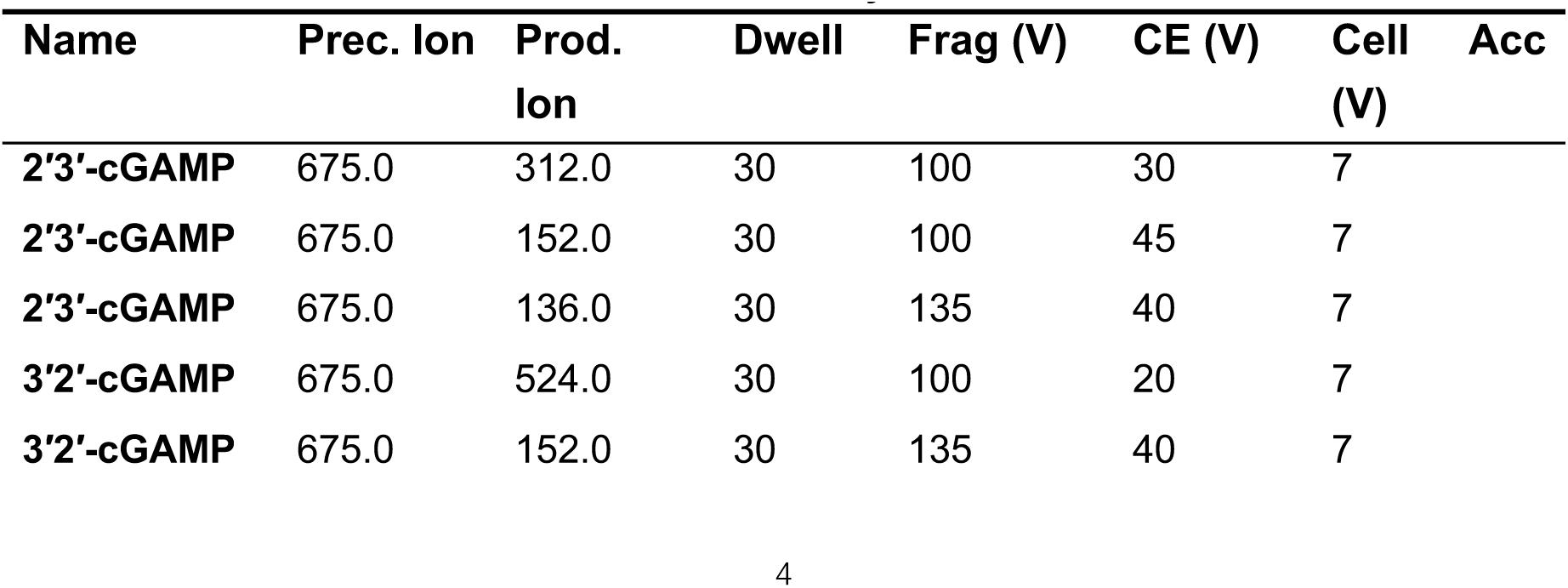
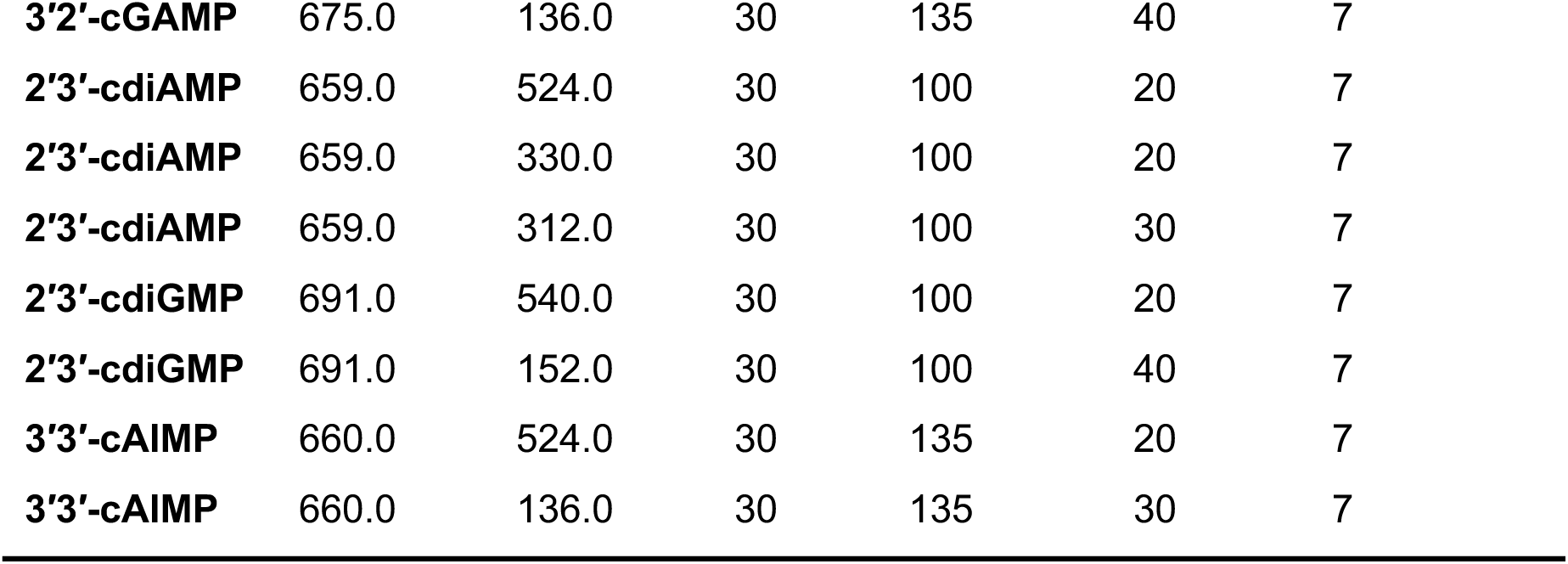
Transition List, Dwell, Frag (V) and CE (V) Cell acc (V) were used to monitor CDNs by MRM.

| **Name** | **Prec. Ion** | **Prod. Ion** | **Dwell** | **Frag (V)** | **CE (V)** | **Cell (V)** | **Acc** |
| --- | --- | --- | --- | --- | --- | --- | --- |
| **2'3'-cGAMP** | 675.0 | 312.0 | 30 | 100 | 30 | 7 |  |
| **2'3'-cGAMP** | 675.0 | 152.0 | 30 | 100 | 45 | 7 |  |
| **2'3'-cGAMP** | 675.0 | 136.0 | 30 | 135 | 40 | 7 |  |
| **3'2'-cGAMP** | 675.0 | 524.0 | 30 | 100 | 20 | 7 |  |
| **3'2'-cGAMP** | 675.0 | 152.0 | 30 | 135 | 40 | 7 |  |

|  |  |  |  |  |  |  |
| --- | --- | --- | --- | --- | --- | --- |
| **3'2'-cGAMP** | 675.0 | 136.0 | 30 | 135 | 40 | 7 |
| **2'3'-cdiAMP** | 659.0 | 524.0 | 30 | 100 | 20 | 7 |
| **2'3'-cdiAMP** | 659.0 | 330.0 | 30 | 100 | 20 | 7 |
| **2'3'-cdiAMP** | 659.0 | 312.0 | 30 | 100 | 30 | 7 |
| **2'3'-cdiGMP** | 691.0 | 540.0 | 30 | 100 | 20 | 7 |
| **2'3'-cdiGMP** | 691.0 | 152.0 | 30 | 100 | 40 | 7 |
| **3'3'-cAIMP** | 660.0 | 524.0 | 30 | 135 | 20 | 7 |
| **3'3'-cAIMP** | 660.0 | 136.0 | 30 | 135 | 30 | 7 |

**Table S4.** HPLC gradient.

|  | **Time** | **A** | **B** | **Flow** |
| --- | --- | --- | --- | --- |
| **1** | 5.00 min | 10.00% | 90.00% | 0.150 mL/min |
| **2** | 5.10 min | 95.00% | 5.00% | 0.500 mL/min |
| **3** | 10.00 min | 95.00% | 5.00% | 0.500 mL/min |
| **4** | 10.20 min | 10.00% | 90.00% | 0.300 mL/min |
| **5** | 15.00 min | 10.00% | 90.00% | 0.300 mL/min |
| **6** | 15.40 min | 10.00% | 90.00% | 0.150 mL/min |
| **7** | 16.00 min | 10.00% | 90.00% | 0.150 mL/min |

**Table S5.** Table S5 The source parameters.

| **Parameter** | **Value (+)** | **Value (-)** |
| --- | --- | --- |
| **Gas temp (°C)** | 350 | 350 |
| Gas Flow (L min<sup>-1</sup>) | 7 | 7 |
| Nebulizer (psi) | 35 | 35 |
| SheathGasHeater | 350 | 350 |
| SheathGasFlow | 12 | 12 |
| Capillary (V) | 2500 | 3500 |
| VCharging | 800 | 500 |

**Table S6.** Transition List, Dwell, Frag (V) and CE (V) Cell acc (V) were used to monitor CDNs by MRM.

| Name | Prec. Ion | Prod. Ion | Dwell | Frag (V) | CE (V) | Cell (V) | Acc |
| --- | --- | --- | --- | --- | --- | --- | --- |
| **2'3'-cGAMP** | 675.0 | 312.0 | 30 | 100 | 30 | 7 |  |
| **2'3'-cGAMP** | 675.0 | 152.0 | 30 | 100 | 45 | 7 |  |
| **2'3'-cGAMP** | 675.0 | 136.0 | 30 | 135 | 40 | 7 |  |
| **3'2'-cGAMP** | 675.0 | 524.0 | 30 | 100 | 20 | 7 |  |
| **3'2'-cGAMP** | 675.0 | 152.0 | 30 | 135 | 40 | 7 |  |
| **3'2'-cGAMP** | 675.0 | 136.0 | 30 | 135 | 40 | 7 |  |
| **2'3'-cdiAMP** | 659.0 | 524.0 | 30 | 100 | 20 | 7 |  |
| **2'3'-cdiAMP** | 659.0 | 330.0 | 30 | 100 | 20 | 7 |  |
| **2'3'-cdiAMP** | 659.0 | 312.0 | 30 | 100 | 30 | 7 |  |
| **2'3'-cdiGMP** | 691.0 | 540.0 | 30 | 100 | 20 | 7 |  |
| **2'3'-cdiGMP** | 691.0 | 152.0 | 30 | 100 | 40 | 7 |  |
| **3'3'-cAIMP** | 660.0 | 524.0 | 30 | 135 | 20 | 7 |  |
| **3'3'-cAIMP** | 660.0 | 136.0 | 30 | 135 | 30 | 7 |  |

