## Supplemental Figures for "A novel virus-induced cyclic dinucleotide, 2′3′-c-di-GMP, mediates STING-dependent antiviral immunity in *Drosophila*"

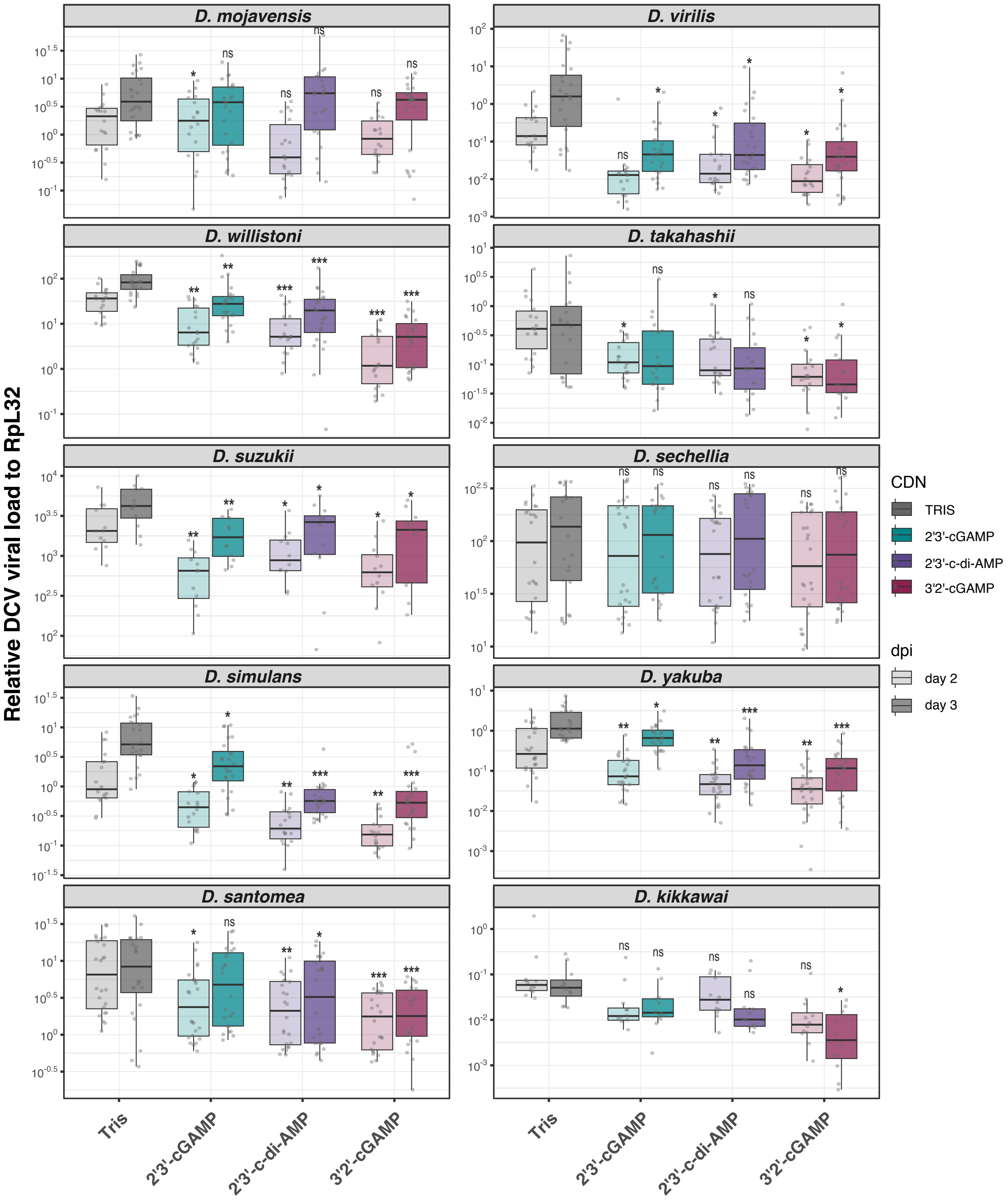


**Supplementary Figure 1: 3′2′-cGAMP induces antiviral immunity in most, but not all, *Drosophila* species**
Relative DCV RNA loads at two or three days post-infection in flies from the indicated ten *Drosophila* species pre-injected with Tris or the indicated CDNs. Data are from at least three independent experiments, each performed in biological triplicates, and shown with boxplot with scatter plot. Data were analysed using pairwise permutation test with FDR method upon CDN-compared to Tris-injection is shown (ns= non significant, *<0.05, **<0.01 and ***<0.001).

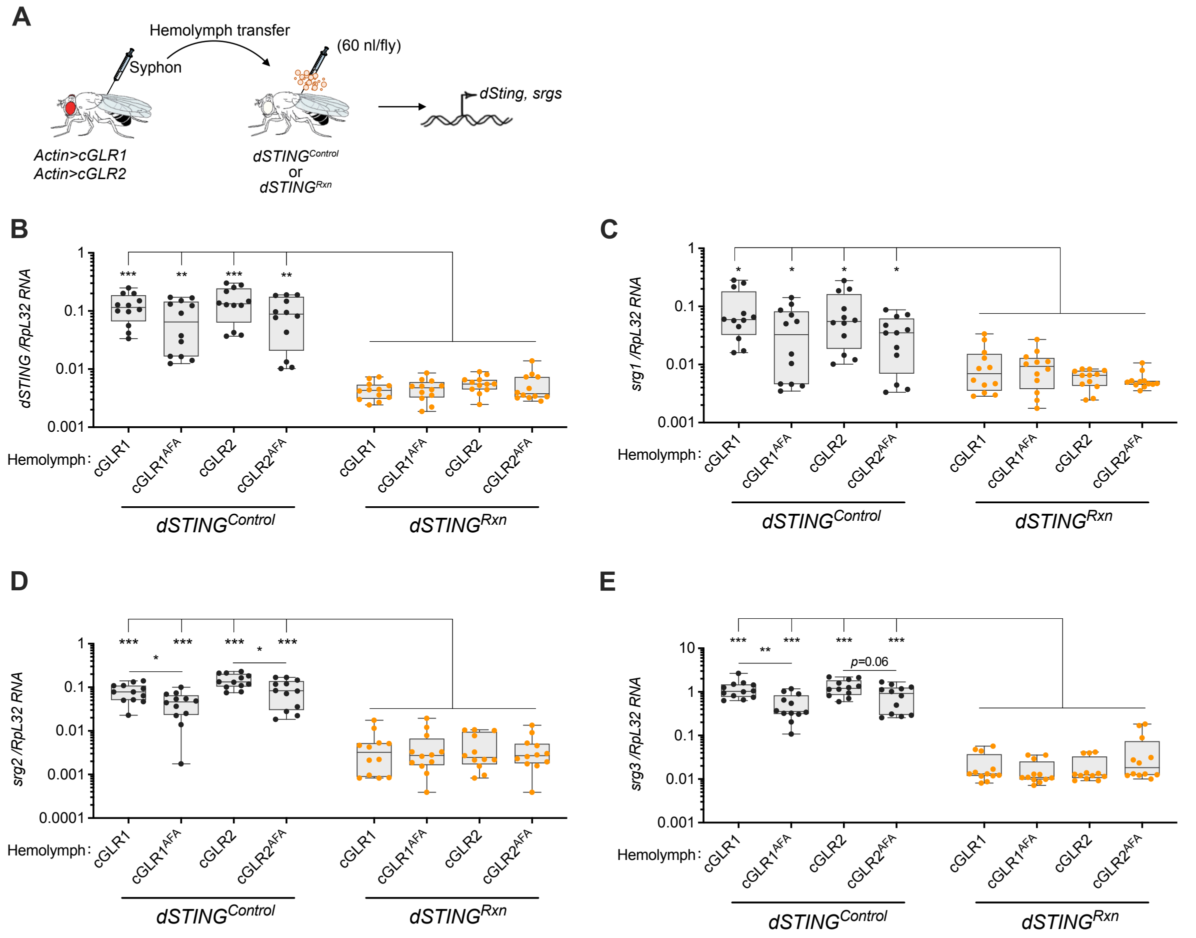


**Supplementary Figure 2: Detection of a STING agonist activity in the hemolymph of cGLRs transgenic flies**

**A.** Hemolymph was extracted from transgenic flies ectopically expressing cGLR1 or cGLR2 or their inactive $cGLR^{AFA}$ versions and injected into naïve control (*$dSTING^{Control}$*) or *dSTING* mutant flies (*$dSTING^{Rxn}$*). Expression of *STING*-regulated genes was monitored to assess activation of the pathway. **B-E.** Relative gene expression of *dSTING* (**B**), *srg1* (**C**), *srg2* (**D**) and *srg3* (**E**) in control or *dSTING* mutant flies 12h after transfer of the hemolymph extracted from flies expressing the indicated transgenes. Data are from four independent experiments, each involving three groups of 6 males flies shown with boxplot with scatter plot. Data were analysed using pairwise permutation test with FDR method (*<0.05, **<0.01 and ***<0.001).

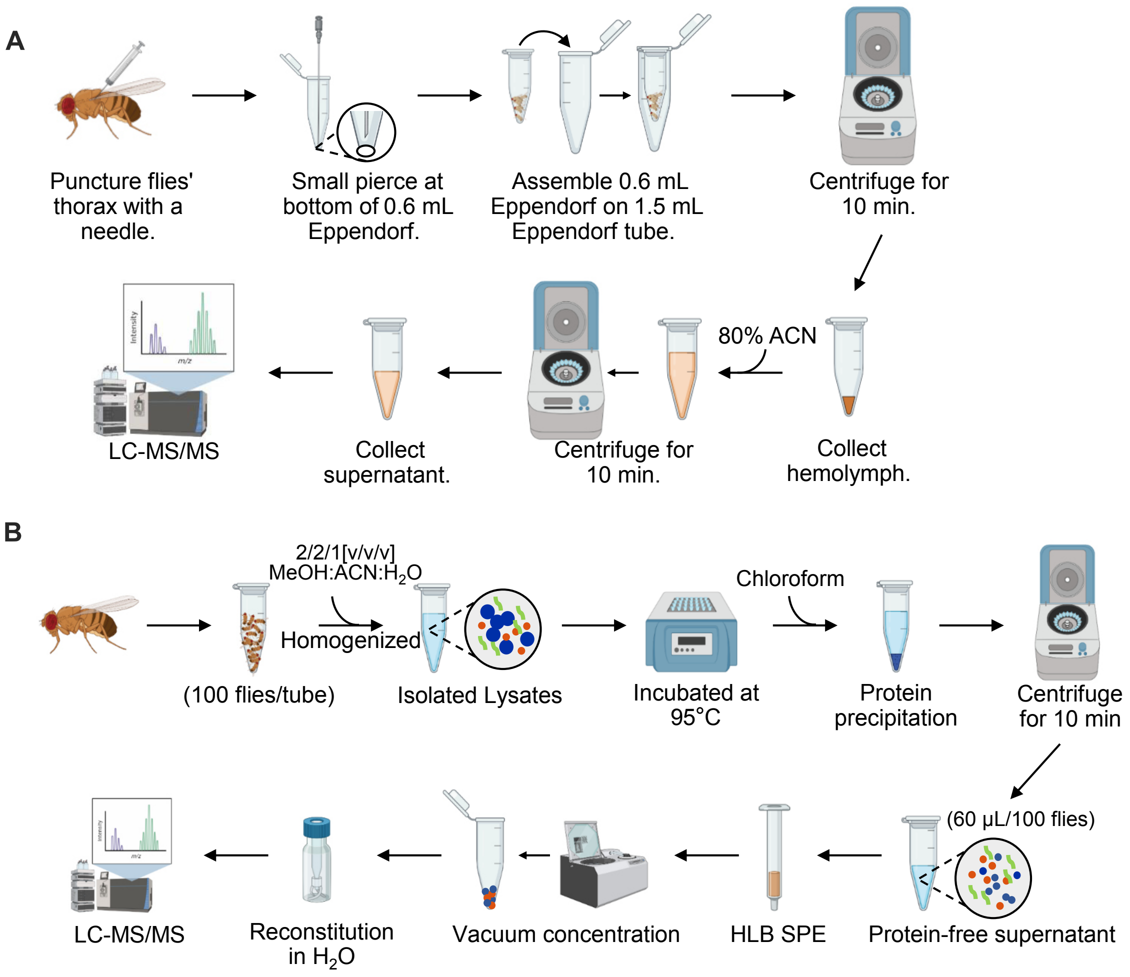


**Supplementary Figure 3: Preparation of extracts from hemolymph or whole-fly lysates for the detection of CDNs using LC-MS analysis**

**A.** For preparation of extracts from hemolymph, punctured flies were collected and transferred into a microcentrifuge tube with a hole in the bottom, which was then placed in a 1.5mL tube and centrifuged to collect the hemolymph. **B.** Whole-fly lysates were incubated at 95°C and extracted with chloroform to eliminate proteins. After centrifugation, the supernatant was applied to a Solid Phase Extraction (SPE) column. The resulting material was vacuum-dried and re-suspended in water for LC-MS/MS analysis.

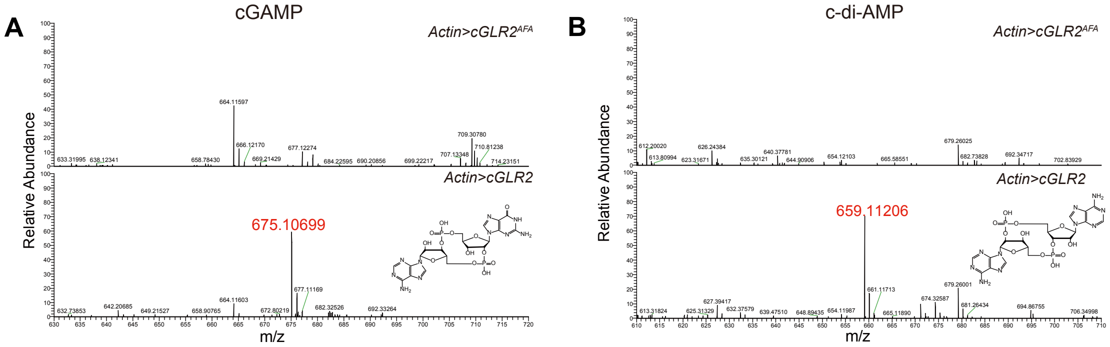


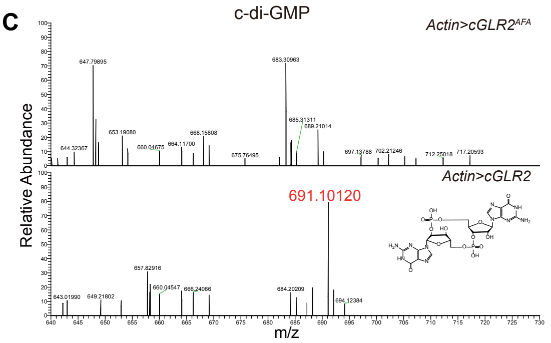


**Supplementary Figure 4: Detection of CDNs from the lysate of flies ectopically expressing cGLR2 or cGLR2AFA using LC-MS analysis**

**A-C.** Full scan mode of LC-MS showing the detection of cGAMP (**A**), c-di-AMP (**B**) and c-di-GMP (**C**) in whole-fly extracts from transgenic flies ectopically expressing cGLR2. **D-F.** Tandem mass spectra after HCD fragmentation of the ion with m/z=675.1072 (**D**), 659.1123 (**E**), 691.1021 (**F**). Arrows indicate the m/z values of the expected fragmentation patterns of 2'3'-cGAMP, 3'2'-cGAMP (**D**), 2'3'-c-di-AMP (**E**) and 2'3'-c-di-GMP (**F**). Data are from one experiment, representative of three independent experiments.

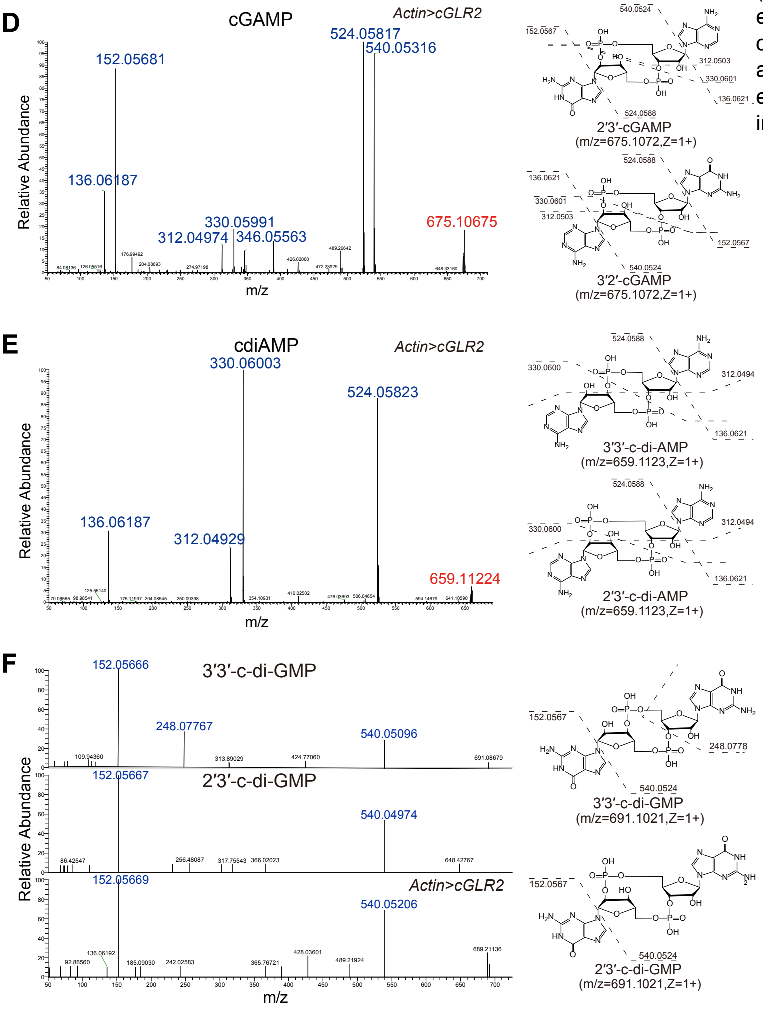

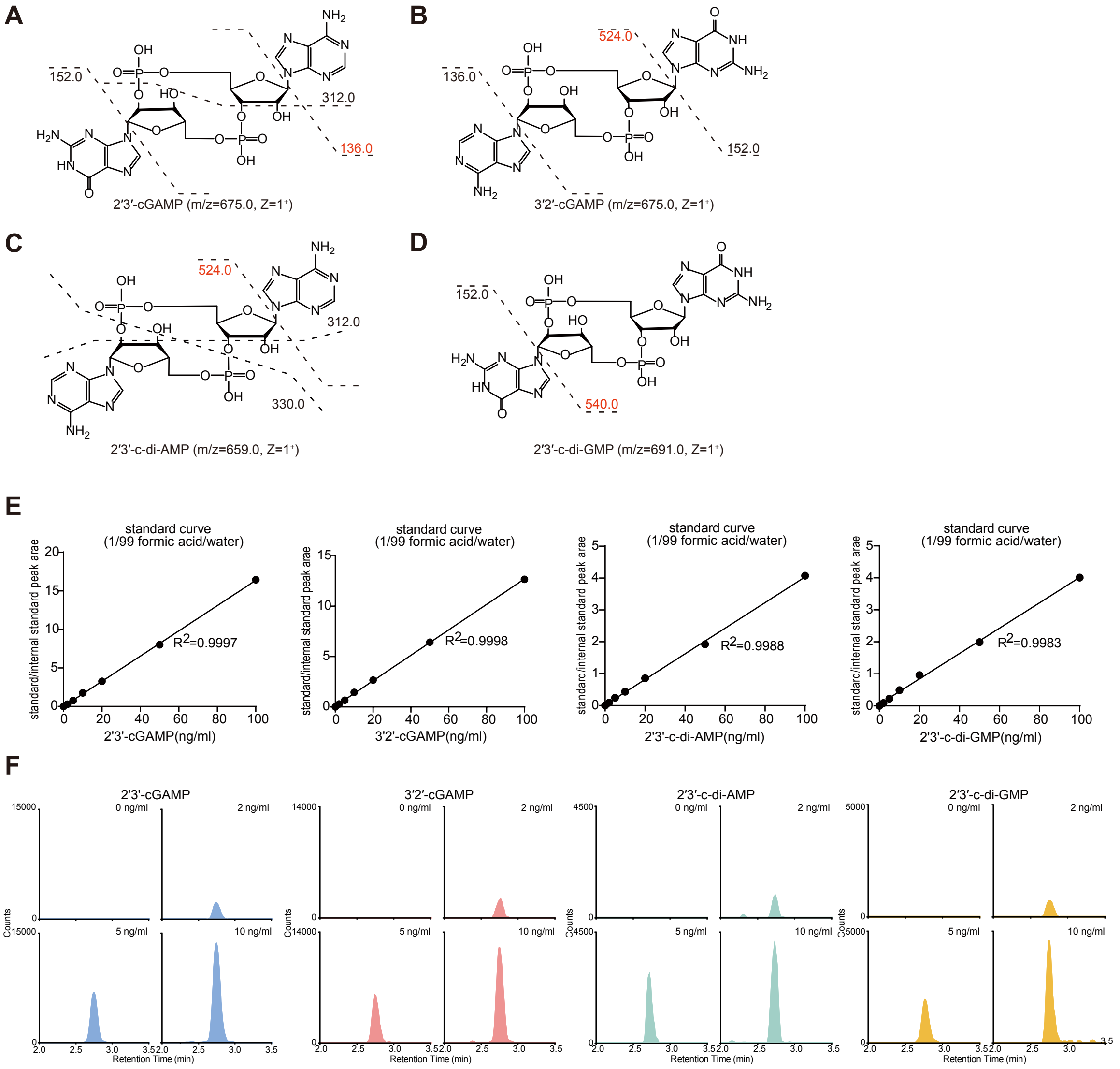


**Supplementary Figure 5: Detection and quantification of CDNs using LC-MS**

**A-D.** Fragmentation pattern of the ions with m/z=675.0 (**A**), 675.0 (**B**), 659.0 (**C**), and 691.0 (**D**). Numbers labelled in red indicate quantifier ions. **E-F.** Standard curves (**E**) and LC traces (**F**) of lowest CDNs standards in 99.9/0.1 acetonitrile/water spiked in directly. The peak area belongs to the quantifier ion of the indicated CDNs (m/z=spectra of quantifier ion). $R^2$ = coefficient of determination, determined after linear least squares regression. Data are from one experiment, representative of 72 independent experiments.

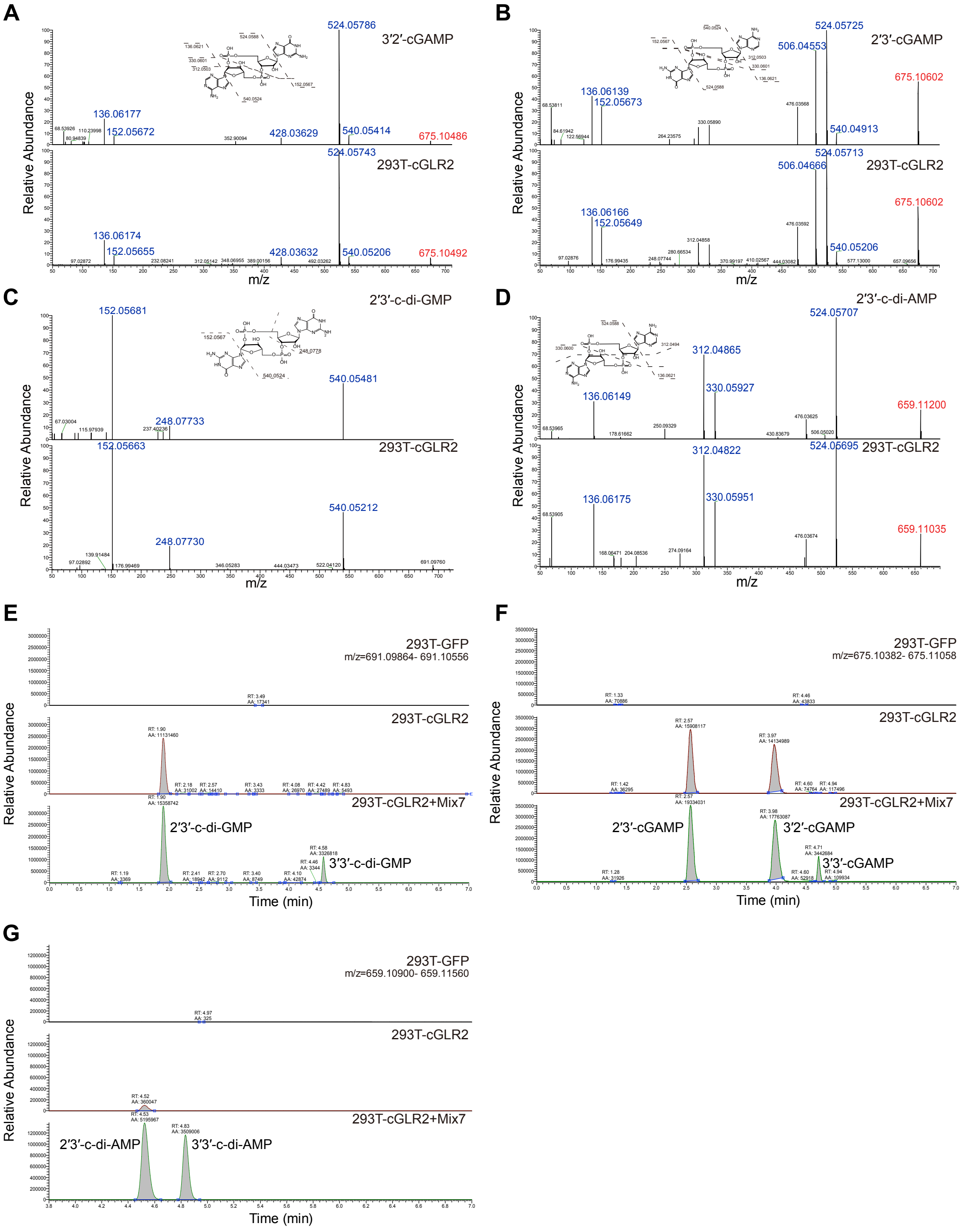


**Supplementary Figure 6: 3′2′-cGAMP, 2′3′-cGAMP, 2′3′-c-di-GMP and 2′3′-c-di-AMP are produced in HEK293T cells expressing cGLR2**

**A-D.** Comparison of tandem mass spectra between extracts from cGLR2 transfected HEK293T cells and chemically synthesized CDNs validate the production of 3′2′-cGAMP (**A**), 2′3′-cGAMP (**B**), 2′3′-c-di-GMP (**C**), 2′3′-c-di-AMP (**D**). **E-G.** Spiking the indicated CDNs in the extracts from cGLR2-transfected HEK293T cells increases the amount of CDNs with the same retention time, confirming the production of 2′3′-c-di-GMP, 2′3′-cGAMP, 3′2′-cGAMP, 2′3′-c-di-AMP. Mix7: a chemical standard mixture of 2′3′-c-di-GMP, 3′3′-c-di-GMP, 2′3′-cGAMP, 3′2′-cGAMP, 3′3′-cGAMP, 2′3′-c-di-AMP, 3′3′-c-di-AMP. Representative of 6 independent experiments.

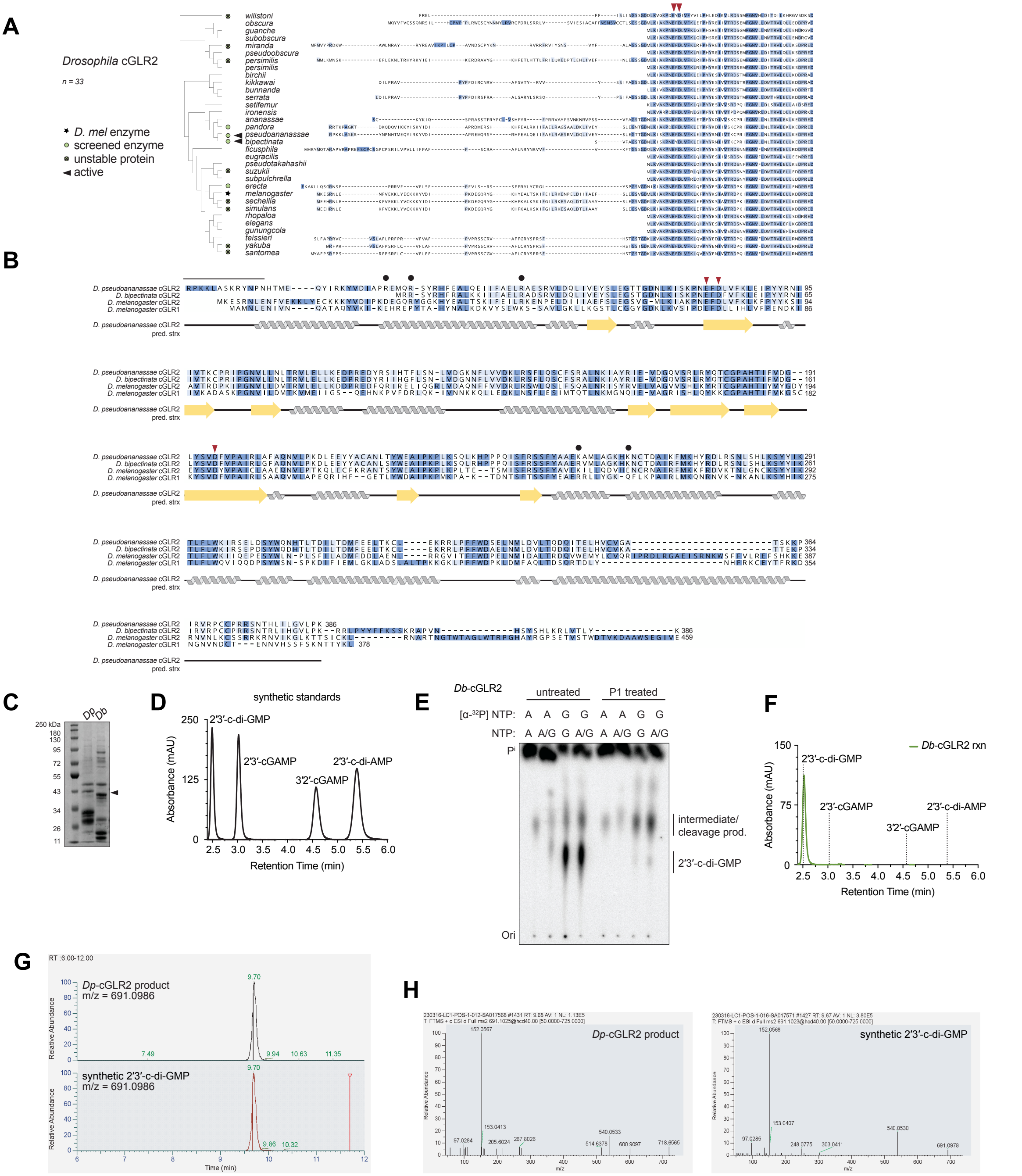


**Supplementary Figure 7: Discovery of 2′3′-c-di-GMP as a nucleotide second messenger product by *Drosophila* cGLR2**
**A.** Sequence alignment (N-terminus shown) and phylogenetic tree of 33 *Drosophila* cGLR2 enzymes. *D. melanogaster* cGLR2 denoted with black star; enzymes selected for recombinant *E. coli* expression denoted with green circle; unstable proteins that could not be purified denoted by black "x"; active cGLR2 enzymes denoted with black triangle; conserved cGLR catalytic residues annotated on sequence alignment with burgundy triangles. **B.** Annotated sequence alignment of relevant cGLR enzymes, shown with predicted secondary structure of *D. pseudoananassae* cGLR2. Residues selected for mutagenesis denoted with black circle; N-terminal amino acids selected for truncation denoted with black line. **C.** SDS-PAGE and Coomassie stain of purified *Dp*-cGLR2 and *Db*-cGLR2 enzymes used in biochemical experiments. **D.** C18 elution profiles of synthetic nucleotide standards. **E.** TLC analysis of α-$^{32}$P labeled *Db*-cGLR2 reaction products and treatment with P1 nuclease. Data are representative of n= 3 independent experiments. **F.** HPLC chromatogram showing C18 elution profile of *Db*-cGLR2 reaction; dashed lined indicates retention times of synthetic nucleotide standards. Data are representative of n= 3 independent experiments. **G, H** The *Dp*-cGLR2 *in vitro* reaction product was analyzed by tandem MS and compared with synthetic 2′3′-c-di-GMP standard. Parent mass ion trace, G, and tandem MS spectra comparison, H validate the chemical identity of the cGLR2 product as 2′3′-c-di-GMP.

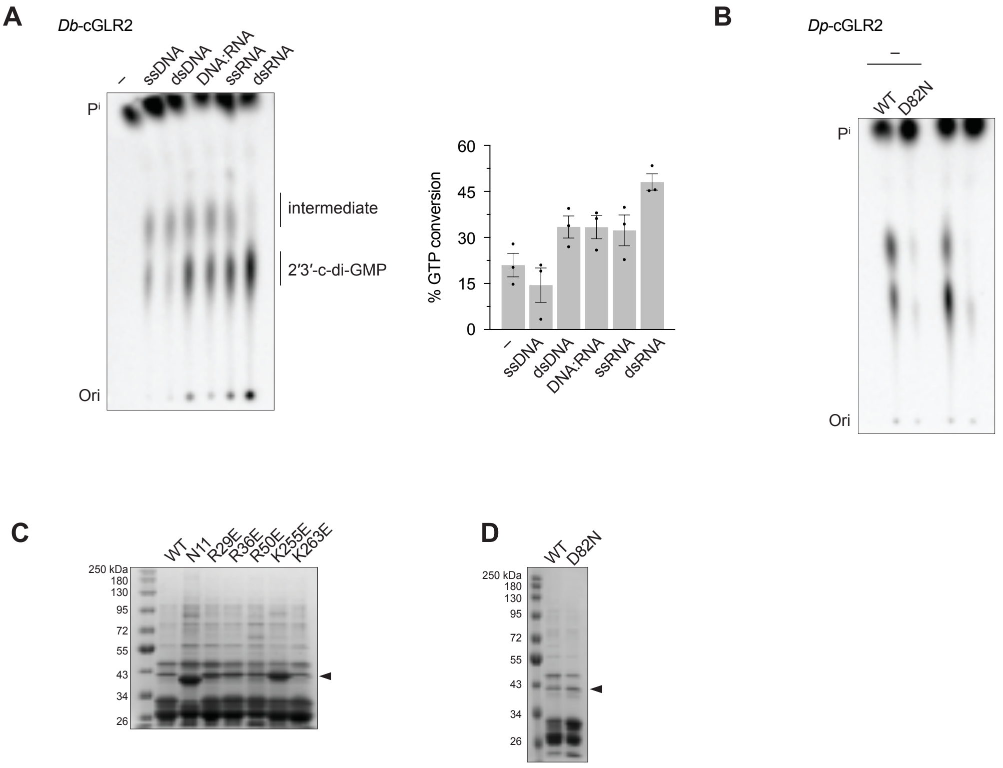


**Supplementary Figure 8: dsRNA binding of Drosophila cGLR2 facilitates its production of 2'3'-c-di-GMP**

**A.** TLC analysis and quantification of GTP conversion to 2′3′-c-di-GMP by *Db*-cGLR2 in the presence of different nucleic acid ligands. Data are mean +/- s.e.m. of n= 3 individual experiments. **B.** TLC analysis of wildtype and D82N mutant *Dp*-cGLR2 reactions. Data are representative of n= 3 independent experiments. **C.** SDS-PAGE and Coomassie stain of purified WT and mutant *Dp*-cGLR2 enzymes used in Fig. 4D. **D.** SDS-PAGE and Coomassie stain of purified WT and mutant *Dp*-cGLR2 enzymes used in Fig. S8B

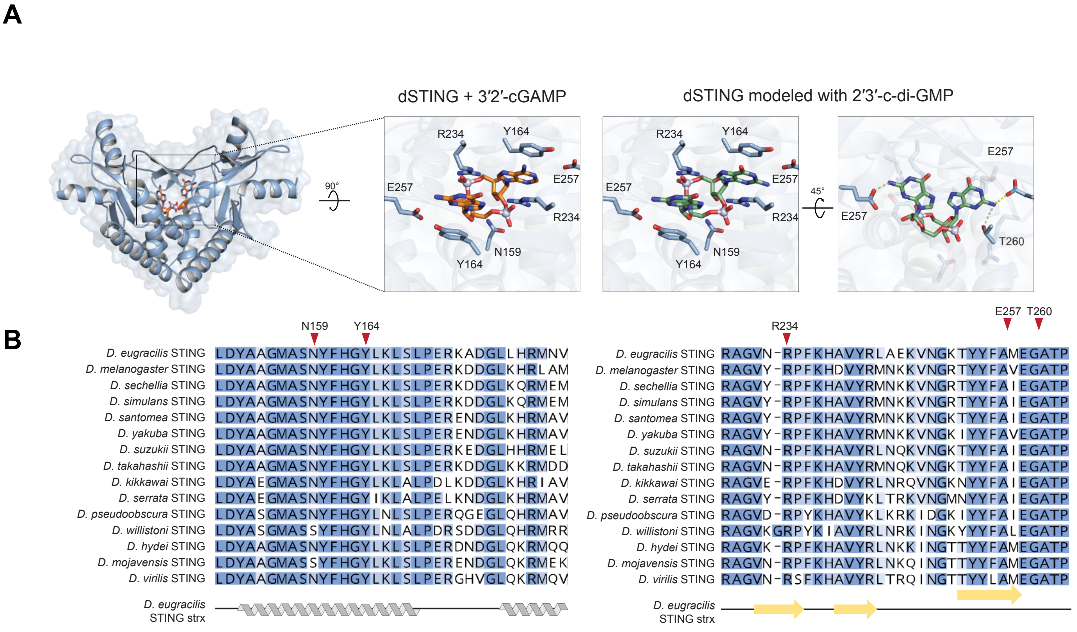


**Supplementary Figure 9: Sequence alignment of STING proteins from relevant *Drosophila* species**

**A.** Crystal structure of *D. eugracilis* STING–3′2′-cGAMP complex (PDB: 7MWZ). Inset (left) highlights important contacts for 3′2′-cGAMP recognition. Insets (right): dSTING with 2′3′-c-di-GMP modeled in the position of 3′2′-cGAMP. In addition to the contacts that control 3′2′-cGAMP recognition, T260 additionally is predicted to contact the N2 position on the guanosine in 2′3′-c-di-GMP which replaces adenosine in 3′2′-cGAMP. **B.** Sequence alignment of STING proteins from relevant *Drosophila* species shown with secondary structure of *D. eugracilis* STING. Critical cyclic dinucleotide binding residues denoted with red triangle and annotated with *D. eugracilis* STING residue number.

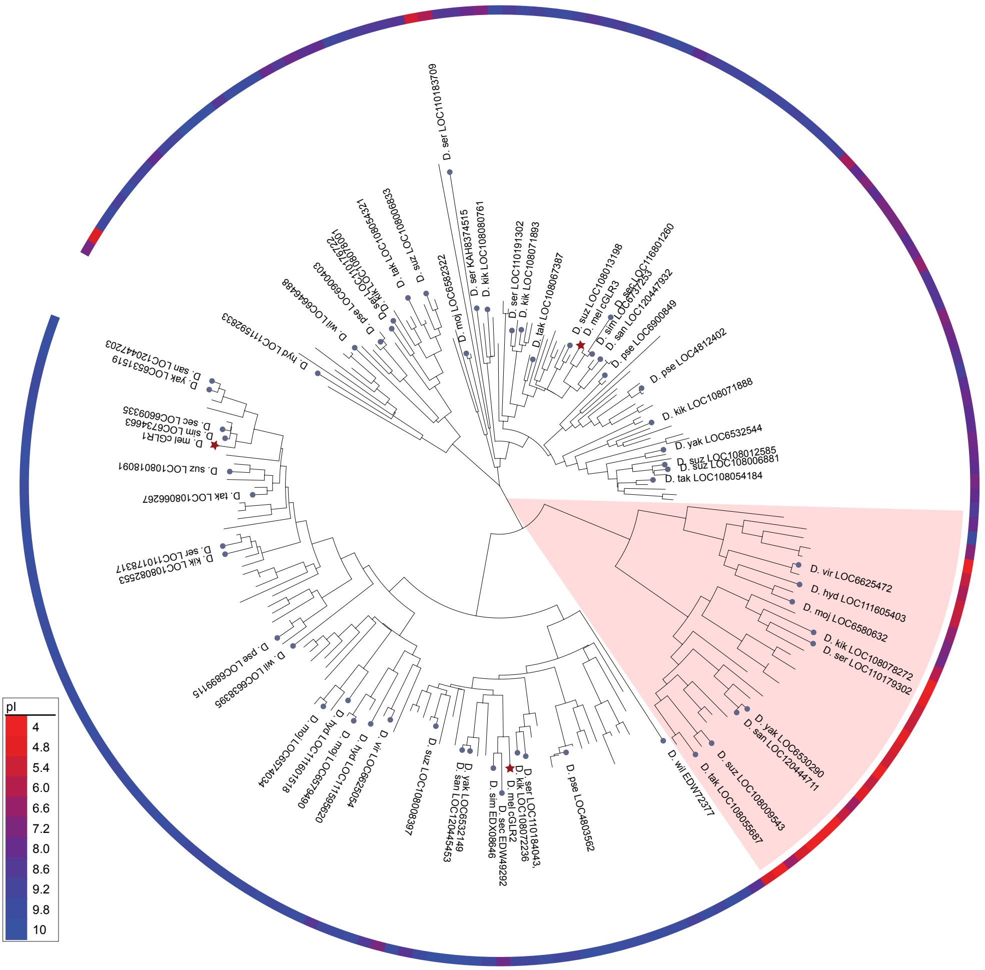


**Supplementary Figure 10: Bioinformatic identification and phylogenetic trees showing all cGLR proteins identified in *Drosophila* genomes**

Isoelectric point (pI) annotated by color gradient where acidic pIs are shown in red and basic pIs are shown in blue.
